# A Peptide Derived from Sorting Nexin 1 Inhibits HPV16 Entry, Retrograde Trafficking, and L2 Membrane Spanning

**DOI:** 10.1101/2024.05.25.595865

**Authors:** Shuaizhi Li, Zachary L. Williamson, Matthew A. Christofferson, Advait Jeevanandam, Samuel K. Campos

## Abstract

High risk human papillomavirus (HPV) infection is responsible for 99% of cervical cancers and 5% of all human cancers worldwide. HPV infection requires the viral genome (vDNA) to gain access to nuclei of basal keratinocytes of epithelium. After virion endocytosis, the minor capsid protein L2 dictates the subcellular retrograde trafficking and nuclear localization of the vDNA during mitosis. Prior work identified a cell-permeable peptide termed SNX1.3, derived from the BAR domain of sorting nexin 1 (SNX1), that potently blocks the retrograde and nuclear trafficking of EGFR in triple negative breast cancer cells. Given the importance of EGFR and retrograde trafficking pathways in HPV16 infection, we set forth to study the effects of SNX1.3 within this context. SNX1.3 inhibited HPV16 infection by both delaying virion endocytosis, as well as potently blocking virion retrograde trafficking and Golgi localization. SNX1.3 had no effect on cell proliferation, nor did it affect post-Golgi trafficking of HPV16. Looking more directly at L2 function, SNX1.3 was found to impair membrane spanning of the minor capsid protein. Future work will focus on mechanistic studies of SNX1.3 inhibition, and the role of EGFR signaling and SNX1-mediated endosomal tubulation, cargo sorting, and retrograde trafficking in HPV infection.

## Introduction

Human papillomaviruses (HPV) are among the most common sexually transmitted pathogens in the United States [1]. More than 220 HPV types have been identified and annotated, with reference genomes available in the PaVE database [2]. Within the genus *Alphapapillomavirus* there are 14 high-risk HPV types, of which HPV16 and HPV18 are the most prevalent [3]. High-risk HPV is responsible for 99% of cervical cancer and just under 5% of all human cancers worldwide [4,5]. HPV16 is alone responsible for roughly 50% of all cervical cancers in women and ∼90% of the HPV-associated oropharyngeal cancers, mostly in men [4,6]. The nonavalent Gardasil-9 vaccine targets high-risk HPV16, HPV18, HPV31, HPV33, HPV45, HPV52, HPV58, and low risk HPV6 and HPV11, but it does not cover all the high-risk HPV types and the high cost prevents people in the low-income region to get access to the vaccine [4,7,8]. HPVs are small non-enveloped DNA viruses, with 72 disulfide-linked pentamers of major capsid protein L1 forming the icosahedral capsid [9]. Variable copies but less than 72 molecules (typically 20-40 molecules) of minor capsid protein L2 are incorporated within the viral particle, complexed with the ∼8kb circular double-stranded viral DNA genome (vDNA) [10,11]. Although only a minor component of the virion, L2 plays an essential role in transporting the vDNA to the host cell nucleus during infection [12].

Successful infection requires HPV16 to gain access to the extracellular matrix (ECM)-rich basement membrane and basal keratinocytes of the differentiated epithelium [13]. At the cellular level, the initial attachment of the HPV16 to keratinocytes is via cell surface and ECM-localized heparin sulfate proteoglycans (HSPGs) [14,15]. Laminin-332 within the ECM has also been identified as an attachment molecule [16,17]. HSPG binding and extracellular furin- and kallikrein-dependent cleavage of the capsid proteins triggers conformational changes within the virion, allow it to engage a secondary entry complex [18–20]. Despite extensive study, the precise nature and composition of this entry complex remains obscure, but it likely contains growth factor receptor tyrosine kinases, integrins, tetraspanins, and annexin A2 [21–23]. Due in part to the time required for entry receptor complex assembly and capsid priming, HPV16 is asynchronously endocytosed by the host cell via macropinocytosis-like mechanisms, which are actin dependent but clathrin- and lipid raft-independent [18,24]. These uptake mechanisms are dependent upon signaling cascades initiated by receptor kinases such as EGFR and Abl2; bursts of EGFR signaling lead to activation of ERK and induction of pit formation while Abl2 kinase regulates maturation of endocytic pits during viral uptake [25]. Efficient trafficking of HPV16 also requires the inhibition of autophagy achieved via another EGFR signaling cascade that the virus uses to prime the cell for viral uptake [26,27]. Downstream of EGFR in this signaling cascade is the phosphoinositol 3-kinase/Akt/mTOR pathway, which can halt autophagy in the cell from activation by HPV16 [27]. EGFR has been known as one of the most important growth factor receptors in HPV entry and has had its role expanded and understood further in recent years [28].

Internalized virions traffic to the endosomal compartment, where the acidic environment in the endosome triggers the further disassembly of the viral capsid allowing for exposure of the L2/vDNA complex [29]. Meanwhile, the γ-secretase protease complex promotes membrane insertion of L2, to allow for spanning and protrusion of L2 through the vesicular membrane [30,31] via an N-terminal transmembrane domain [32]. Stable membrane spanning of L2 results in the cytosolic exposure of the C-terminal ∼400 residues containing binding regions for cellular cargo sorting factors including SNX17, SNX27, and the heterotrimeric retromer complex [33–37]. Binding of these cytosolic factors to the cytosolic portion of transmembrane L2 facilitates efficient retrograde transport of the vesicle-bound vDNA to the *trans*-Golgi network [38], an obligate step of infection. Post-Golgi trafficking requires the onset of mitosis [39,40]. During this process, L2’s chromosome-binding domain tethers the vDNA to host chromosomes to achieve nuclear delivery of L2/vDNA upon onset of the next G1 phase [41]. After mitosis, L2/vDNA recruits PML body components for efficient viral gene expression within the basal keratinocytes [42–44]. Once initial infection is established, episomal HPV16 vDNA will remain within these at a low copy number [13,45]. As the infected basal cells differentiate, expression of early viral genes, vDNA amplification, expression of late viral genes L1 and L2, and virion packaging and assembly will be coordinately achieved [46].

Retrograde trafficking of internalized L2/vDNA is a required step for HPV infection. The SNX-BAR subfamily of sorting nexins are key players in endocytosis, endosomal sorting, and vesicle trafficking. This family includes SNX1, SNX2, SNX4, SNX5, SNX6, SNX7, SNX8, SNX9, SNX18, SNX30, SNX32 and SNX33 [47]. A subgroup of the SNX-BAR family members, comprising SNX1, SNX2, SNX5, and SNX6 support sorting and retrograde transport of cargo from endosomes to the the Golgi [47,48]. SNX-BAR family proteins contain two major domains; the N-terminal PX domain that bind to specific phosphatidylinositol lipids in cellular membranes and a curved BAR domain that promotes protein-protein interactions and membrane remodeling [47,49]. Structurally, the curved BAR domains can form homo- and heterodimers with other SNX-BAR proteins to augment PX domain-dependent binding of the SNX-BAR protein to curved membranes, thereby supporting the tubulation of vesicles and the sorting of cargo into these tubules for transport. Overexpression of SNX1 can result in BAR domain-dependent bending and extensive tubulation of endosomal membranes. In certain contexts, overexpression of SNX1 also promotes the interaction between the SNX1 BAR domain and EGFR, causing lysosomal degradation of the EGFR, a receptor that has been shown important for HPV infection [50,51]. SNX2, which has a different vesicular distribution pattern than SNX1, and less association with retromer compared with SNX1, has also been shown to promote tubulation and regulate EGFR lysosomal sorting [48,51,52]. Recent studies with cells genetically devoid of retromer components Vps35 and Vps29 have revealed direct binding and sorting of specific cargo receptors by SNX1/5 and SNX2/6 heterodimers, independently of the retromer complex [53–55]. These complexes, which facilitate tubular-based endosomal sorting of specific transmembrane cargo proteins, have been termed “ESCPE-1” and are comprised of SNX1/5 and SNX2/6 heterodimers [53,56].

Recently our colleagues [57] described a peptide therapeutic that kills epidermal growth factor receptor (EGFR)-dependent triple negative breast cancer cells (TNBC) *in vitro* and in mouse models. The peptide is named SNX1.3 and works by potently inhibiting the retrograde nuclear trafficking of EGFR, a pathway crucial for TNBC survival [57]. The peptide is derived from residues 490-515 of the BAR domain of SNX1 (Fig 1A) and is fused to a TAT-derived cell penetrating peptide called PTD4 [58] for efficient cellular uptake. This alpha helical region of SNX1 lies at the interface of SNX1 BAR domain homo- and heterodimers (Fig 1A) and is known to directly bind to the EGFR kinase domain to alter signaling, trafficking and degradation [51,57]. Given that SNX1 is involved in retrograde trafficking of EGFR and other receptor cargos, and that the SNX1.3 peptide alters EGFR retrograde trafficking in TNBC cells [57], and the role of EGFR in HPV16 infection, we sought to determine the effects of SNX1.3 on HPV infection and trafficking.

**Fig 1.**
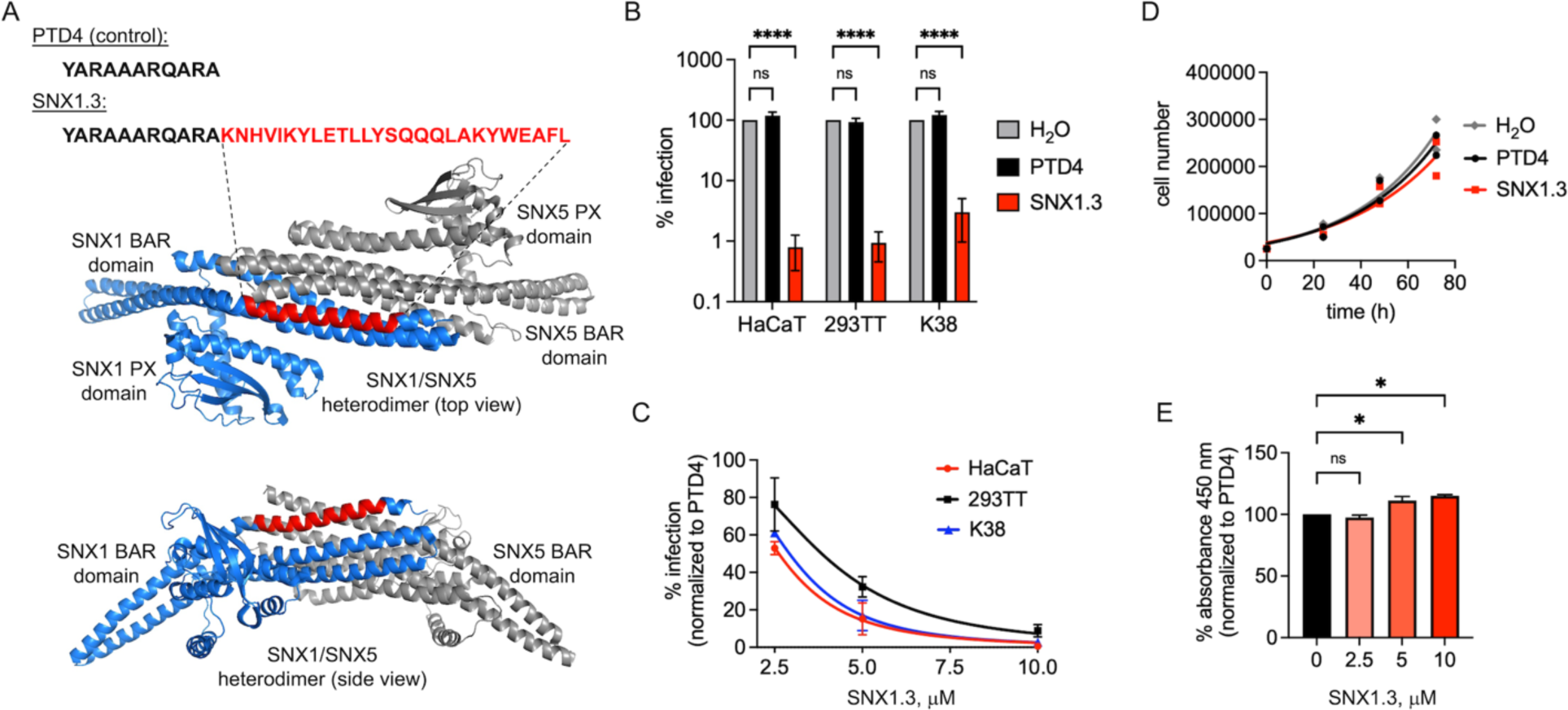
Inhibition of HPV16 infection by SNX1.3 peptide. **(A)** Sequences of the control PTD4 cell penetrating peptide and the SNX1.3 peptide. SNX1.3 is derived from residues 490-515 of the SNX1 BAR domain. This portion is highlighted (red) within the context of the SNX1/SNX5 heterodimer (PDB 8AFZ), at the interface of the BAR domains. **(B)** Infectivity of firefly luciferase expressing HPV16 in multiple cell types in the presence of 10 μM PTD4 or SNX1.3 peptide, compared to H_2_O vehicle control. **(C)** Titration of peptides showing dose-dependent inhibition of HPV16 infection in multiple cell types. For **(B, C)**, the graphs show mean infectivity ± SEM. For **(C)** all groups were statistically significant p<0.05 except for PTD4 vs SNX1.3 in 293TT. **(D)** Growth curves of HaCaT cells ± 10 μM PTD4 or SNX1.3 peptide, compared to H_2_O vehicle control. No statistical significance was observed by between groups at any particular timepoint. **(E)** WST-1 assay for HaCaT cell viability ± PTD4 or SNX1.3 peptide. Assay performed 48h post seeding with peptides. Graph shows mean percent 450nm absorbance ± SEM. All statistics were determined by 2-way ANOVA with Tukey’s multiple comparison. *p<0.05, ****p<0.0001.

Here, we observed strong and specific inhibition of HPV infection in multiple cell types treated with SNX1.3 peptide. Unlike nuclear EGFR-dependent TNBC, SNX1.3 does not affect keratinocyte viability or proliferation. Analysis of subcellular HPV trafficking suggests SNX1.3 treatment delayed viral entry and caused a strong defect in viral retrograde trafficking, attributable to the inhibition of L2 membrane spanning. These data suggest a multi-step inhibition of the HPV infection by SNX1.3 peptide and underscore the need for further mechanistic studies on interactions of the SNX1.3 peptide with cellular targets and studies of SNX-BAR proteins in both viral endocytosis and retrograde sorting/trafficking.

## Materials and Methods

### Tissue culture

Immortalized human HaCaT keratinocytes [46] and murine K38 keratinocytes [59] were cultured in high glucose DMEM media (GIBCO 11965-092) supplied with penicillin and streptomycin (GIBCO 15140-122) and 10% fetal bovine serum (GIBCO A3160602). HaCaT GFP-BAP cells, which stably express cytosolic GFP-BAP [40], were cultured in the the parental HaCaT media supplemented with 0.2ug/ml of puromycin (Corning 61-385-RA) to maintain GFP-BAP expression. 293TT cells were used for generating HPV16 pseudoviruss were cultured in high-glucose DMEM media supplied with 10% bovine growth serum (BGS, HyClone SH30541.03) and 165ug/ml hygromycin B (Invitrogen 10687010) for selection. All cell lines were cultured in 5% CO_2_ tissue culture incubator at 37°C. Cells were passaged when their confluence was around 80-90%. 0.05% trypsin-EDTA (GIBCO® 25300-054) was used for trypsinizing the cells. For cell counting 10μl 1:1 mixed suspended cells and trypan blue solution, 0.4% (w/v) in PBS (Corning 25-900-CI) were applied to a hemocytometer for manual cell counting.

### Peptides

All peptides used in this project were generated by GenScript. Peptide stocks were dissolved in ultrapure water at 1mM final concentration. The sequences are as follows; PTD4: YARAAARQARA-acid, SNX1.3: YARAAARQARAKNHVIKYLETLLYSQQQLAKYWEAFL-acid, SNX2.1: YARAAARQARAKTVIIKYLESLVQTQQQLIKYWEAFL-acid.

### HPV16 PsVs production

293TT cells were cultured in 10cm dishes. At the confluence around 50%, cells were co-transfected with 15ug/plate of pGL3, a luciferase-expressing plasmid with SV40 origin of replication, and 15ug/plate of pXuLL-based plasmids (which express HPV16 L1 and L2) using Ca_3_(PO_4_)_2_ transfection methods. The L2 packaged within these preps are L2-3xFTHA, an HA-tagged form of L2 (L2-HA) previously shown to be infectious [60]. Plasmids were diluted into ultrapure water and 2M CaCl_2_ solution, then mixed with 2xHBS (HEPES, buffered saline, 50mM HEPES, 280mM NaCl, 1.5mM Na_2_PO_4_)) to generate DNA-Ca_3_(PO_4_)_2_ crystal solution. Then it was incubated at room temperature for 30m and added to the cells with gentle mixing. Fresh DMEM media containing 10% BGS was changed the morning after transfection. 293TT cells were harvested at 48h post-transfection. Cells were resuspended in PBS supplied with 9.5mM MgCl_2_. and lysed by 0.35% detergent Brij58. (NH_4_)_2_SO_4_, pH=9.0 was added to 25mM to provide a basic environment to promote L1 disulfide formation and viral capsid maturation [61]. Benzonase nuclease (Sigma E1014) was added to 0.3% final concentration and Plasmid-Safe ATP-dependent DNase (Epicentre E3105K) was added to 20 U/ml final concentration to digest any free unpackaged DNA. Lysates were incubated at 37°C, 5% CO_2_ in the tissue culture incubators for 20h, then ice chilled samples were supplemented with 0.17 volume of 5M NaCl. Lysates were frozen and thawed at -80°C and 37°C to further lyse the cell nuclei and release the viral particles. The samples were centrifuged at 4°C at 10,000 x g for 10m. Pellets from this spin were resuspended in viral storage buffer (VSB, 25mM HEPES with 500mM NaCl and 1mM MgCl2, pH=7.5), and spun again. The two supernatants were pooled and loaded onto the top of the CsCl gradient made by 4ml heavy (1.4g/ml, in VSB) CsCl and 4ml light (1.25g/ml, in VSB) CsCl. Samples were centrifuged at 49,400 x g at 16°C for 18h by using Beckman SW41 Ti rotor/buckets. A visible, blueish-colored viral band appeared near the interface between light and heavy CsCl after centrifugation. 18-gauge needles and 3ml syringes were used for puncturing the side of the centrifuge tube to collect viral bands. 100 kDa MWCO vivaspin concentrator units (Sartorius VS04T42) equilibrated with VSB were used for washing and purifying the collected virus band. After the final wash, the virus was concentrated to 100-200ul and transferred into siliconized tubes for storage at - 80°C. OD_260_ of DNA concentration of purified virus were measured by spectrophotometer to calculate the concentration of physical viral particles. As HPV16 does not have a specific vDNA packaging signal, the content of the pGL3 in the purified viruses was measured by SYBR green qPCR (Thermo Fisher K0252). The serial dilution of the standard curve was used to determine the amount of pGL3 in virus samples, the primers qLuc2-A (ACGATTTTGTGCCAGAGTCC), and qLuc2-B (TATGAGGCAGAGCGACACC) were used for amplifying a portion of the firefly luciferase gene on pGL3. Once the pGL3 concentration was measured, the capsid/pGL3 ratios were determined. Physical titers (viral particles, or ng L1/unit volume) were used for all experiments looking at intracellular levels of virus while pGL3 titers (pGL3 molecules/unit volume) were used for all infection experiments.

### Cell proliferation and cytotoxicity assays

For growth curve assay, 25,000 HaCaTs were seeded in 6 well plates along with 10μM of peptide (PTD4 or SNX1.3) or water control, then after 24h, 48h, and 72h, cells from each group were trypsinized and manually counted using a hemacytometer. For WST-1 assay, HaCaTs were plated at 6,000 cells/well in 96 well plates, in 100μl DMEM supplied with 10% FBS. Once the cells were attached to the bottom, 10μM, 5μM,2.5μM peptides were added to the cell cultures. Cells were incubated at 37°C and 5% CO_2_ in the presence of 2.5μM, 5μM, and 10μM peptides for 48h. Then, 10μl /well cell proliferation reagent WST-1 (Roche 05015944001) was added, and cells were incubated at 37°C, 5% CO_2_ for another 4h. Plates were thoroughly shaken for 1m, and the absorbance of formazan product was measured using a DTX800 Multimode plate reader (Beckman Coulter) at 450nm. The cell-free media only group served as the negative control for background subtraction.

### HPV16 PsV infection assay

50,000 cells/well HaCaTs, HaCaT-GFP-BAP, mouse keratinocyte K38 or 80,000 cells/well 293TT cells were seeded in 24 well plates the day before infection. Cells were infected with HPV16 at 2x10^8^ viral genome equivalents (pGL3 copies) per well under the presence of 2.5μM, 5μM, and 10μM peptide. At 24h post-infection, cells were lysed in 100μl reporter lysis buffer (RLB, Promega E397A). 100μl luciferase assay reagent (Promega E1483) was added into 20μl cell lysate and luciferase activity was measured using a DTX800 Multimode plate reader (Beckman Coulter). Western blots and GAPDH immunostaining were performed by using the remaining lysate. GAPDH bands were quantified by densitometry using ImageJ software [62] and used to normalize the luciferase data.

### SDS-PAGE and western blotting

For denaturing/reducing polyacrylamide gel electrophoresis (PAGE), samples were lysed in RIPA lysis buffer (50mM Tris-HCl pH = 8.0, 150mM NaCl, 1% Triton X-100, 0.5% Na-deoxycholate, 0.5% SDS, supplemented with 1% PMSF (Sigma 78830) and 1% protease inhibitor cocktail (Sigma P8340), combined with 20% total volume of denaturing/reducing SDS-PAGE buffer. Samples were then heated to 95°C for 5 minutes before separation on 10% or 12.5% polyacrylamide gels and run at 110V in TGS running buffer (25mM Tris, 192mM glycine, 0.1% SDS, pH=8.6). Samples were transferred onto nitrocellulose membranes under 300mA for 75m by using western transfer buffer (25mM Tris, 192mM glycine) supplemented with 10% methanol. Nitrocellulose membranes were then blocked with 5% non-fat milk in TBST (20mM Tris, 150mM NaCl, 0.1% Tween-20, pH = 7.5) at 4°C overnight. For the denaturing/nonreducing PAGE, samples lysed by RIPA lysis buffer supplemented with 1% PMSF, 1% proteinase inhibitor, and 2mM N-ethylmaleimide (NEM, Sigma E1271) were mixed with 20% total volume of denaturing /non-reducing SDS-PAGE buffer and then incubated at room temperature for 10m before PAGE gel separation. RLB lysed samples were added with 20% total volume of denaturing/reducing SDS-PAGE buffer and incubated at 95°C for 5 mins before SDS-PAGE separation. For translocation experiments, after the transfer step, Nitrocellulose membranes were blocked in Odyssey Blocking Buffer PBS (LI-COR) at 4°C overnight. Antibodies used in western blot are listed in table 2. Blots were imaged on the Licor Odyssey Infrared Imaging System.

### Immunofluorescence staining and confocal microscopy

100,000 cells/well HaCaTs cells were plated on glass coverslips in 6 well plates. Cells were infected with 1μg L1/ml of wt HPV16 PsVs with either 10μM PTD4 or SNX1.3 peptide. For some experiments, NH_4_Cl was added to the infected cells to final concentration of 20mM. For Golgi trafficking experiments, aphidicolin (Santa Cruz 38966-21-1) was added to a final concentration of 6μM. At 2h, 4h, 18h, or 24h post-infection, cells were washed with high pH PBS (pH=10.6-10.8) for 2.5m to remove all the surface-bound viruses, then fixed with 2% paraformaldehyde (pH = 7.2-7.4, Fisher Scientific 30525-89-4) for 15m, permeabilized by 0.25% Triton X-100 (Fisher Scientific 9002-93-1) for 10m and blocked overnight by using blocking solution (PBS plus 4% bovine serum albumin, fraction V, Fisher Scientific 9002-46-8, supplemented with 1% goat serum). Cells were incubated with primary antibody at room temperature for 1h, followed by 1h room temperature incubation of secondary antibody. All antibodies were prepared in PBS containing 20% blocking solution. Prolong diamond anti-fade mounting medium with DAPI (Life Technologies P36971) were used for mounting coverslips to the glass slide. Confocal microscopy was performed by the Zeiss LSM880 system with 405nm, 488nm, and 543nm lasers. Samples were examined using a 63x objective, and Z-stacks with a 0.35μm depth per plane were taken of each image. Images were processed with Zen Blue software and ImageJ software. The Manders’ overlap coefficients were determined using the JACoP plugin on ImageJ under the thresholds that were manually set below saturation [62]. The data were plotted with GraphPad Prism v10 software. Antibodies used in immunofluorescence are listed in Table 1.

**Table 1.**
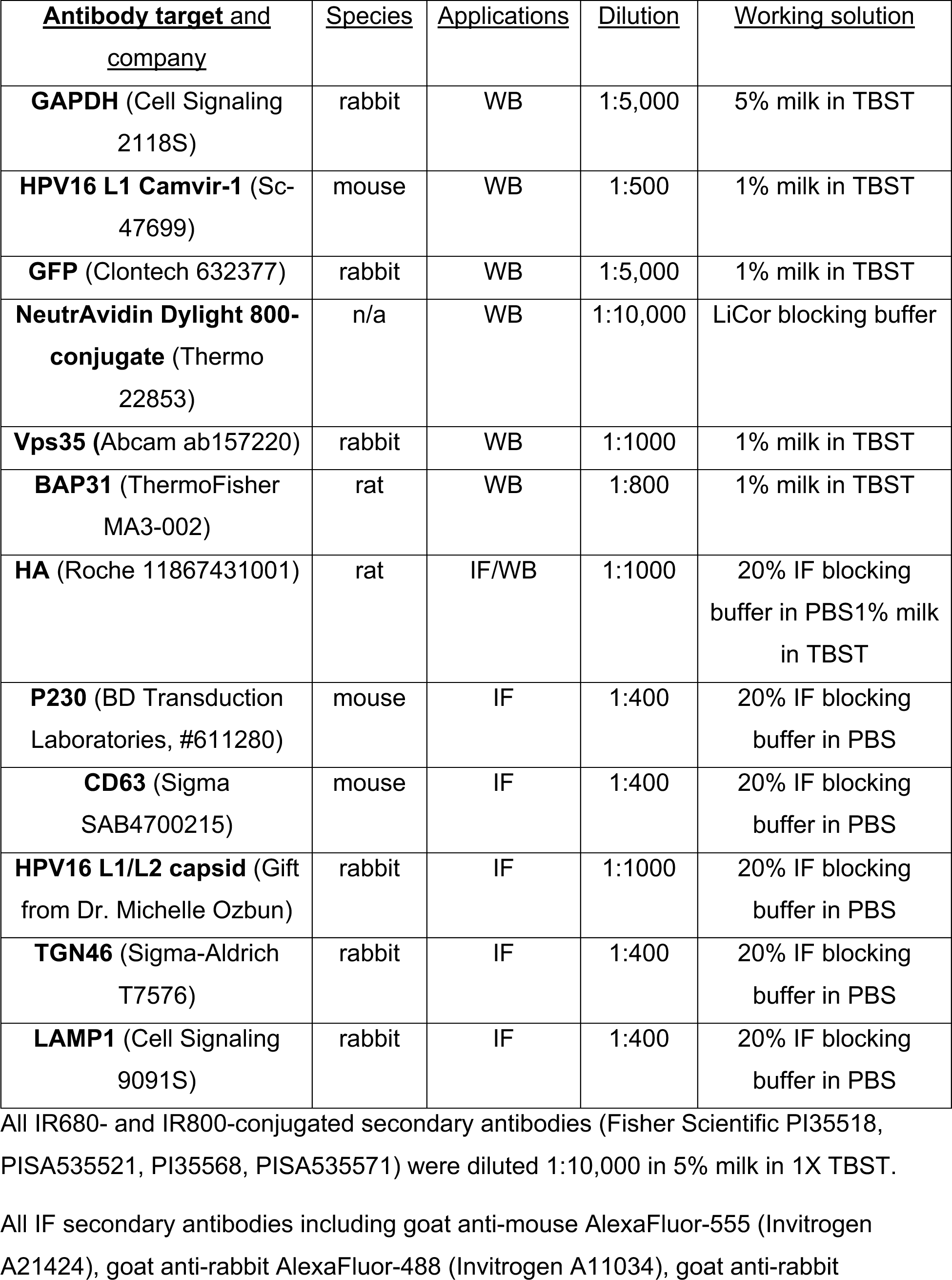
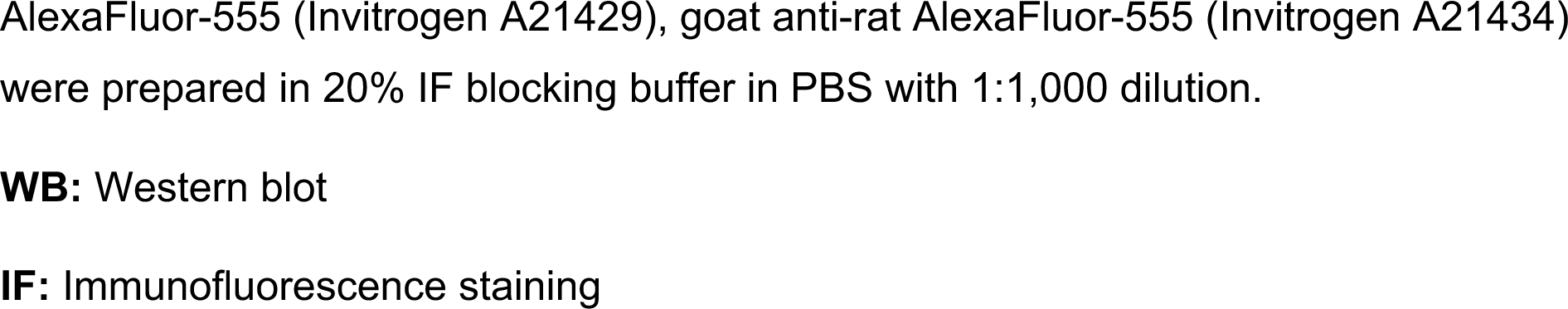
Antibodies used in this work.

| <u>Antibody target and company</u> | <u>Species</u> | <u>Applications</u> | <u>Dilution</u> | <u>Working solution</u> |
| --- | --- | --- | --- | --- |
| **GAPDH** (Cell Signaling 2118S) | rabbit | WB | 1:5,000 | 5% milk in TBST |
| **HPV16 L1 Camvir-1** (Sc-47699) | mouse | WB | 1:500 | 1% milk in TBST |
| **GFP** (Clontech 632377) | rabbit | WB | 1:5,000 | 1% milk in TBST |
| **NeutrAvidin Dylight 800-conjugate** (Thermo 22853) | n/a | WB | 1:10,000 | LiCor blocking buffer |
| **Vps35** (Abcam ab157220) | rabbit | WB | 1:1000 | 1% milk in TBST |
| **BAP31** (ThermoFisher MA3-002) | rat | WB | 1:800 | 1% milk in TBST |
| **HA** (Roche 11867431001) | rat | IF/WB | 1:1000 | 20% IF blocking buffer in PBS<br>1% milk in TBST |
| **P230** (BD Transduction Laboratories, #611280) | mouse | IF | 1:400 | 20% IF blocking buffer in PBS |
| **CD63** (Sigma SAB4700215) | mouse | IF | 1:400 | 20% IF blocking buffer in PBS |
| **HPV16 L1/L2 capsid** (Gift from Dr. Michelle Ozbun) | rabbit | IF | 1:1000 | 20% IF blocking buffer in PBS |
| **TGN46** (Sigma-Aldrich T7576) | rabbit | IF | 1:400 | 20% IF blocking buffer in PBS |
| **LAMP1** (Cell Signaling 9091S) | rabbit | IF | 1:400 | 20% IF blocking buffer in PBS |

All IR680- and IR800-conjugated secondary antibodies (Fisher Scientific PI35518, PISA535521, PI35568, PISA535571) were diluted 1:10,000 in 5% milk in 1X TBST.
All IF secondary antibodies including goat anti-mouse AlexaFluor-555 (Invitrogen A21424), goat anti-rabbit AlexaFluor-488 (Invitrogen A11034), goat anti-rabbit
AlexaFluor-555 (Invitrogen A21429), goat anti-rat AlexaFluor-555 (Invitrogen A21434) were prepared in 20% IF blocking buffer in PBS with 1:1,000 dilution.
**WB:** Western blot
**IF:** Immunofluorescence staining

### Image analysis for entry experiments

A protocol for measurement of total cell intensity using ImageJ [63] was used to quantify the viral entry micrographs. Micrograph data were imported in ImageJ and the selection tool was used to segment the border of single cells of the maximum intensity composites for measurements. After the cells are segmented, the area, mean gray value, and integrated density values were measured. A background region devoid of cells was selected and analyzed from each field of view as well. Once all the measurements were collected, the corrected total capsid channel fluorescence for each segmented region was calculated as integrated density – (area of selected cell – mean fluorescence of background readings). This process was manually done for all cells in each field of view. The majority of the segmented regions contained a single cell, but several contained doublets of cells that were difficult to individually segment. The corrected total intensity of the capsid signal was then calculated and plotted between PTD4 treated and SNX1.3 treated cells using GraphPad Prism v10.

### Viral binding and entry assays

Cells were chilled on ice for 20m before infecting with 1μg L1/ml of HPV16 under the presence of 10μM of peptide in cold DMEM media supplemented with 10% FBS. Plates were then stayed on the ice for 1h to allow surface binding of the viral particle. For the binding sample, cells were first washed with cold PBS (pH = 7.4) to remove all unbound viruses. Then the control groups were washed with cold high-pH PBS (pH = 10.75) to remove the surface-bound virus. Samples were then collected for non-reducing SDS-PAGE. For the entry sample, after 1h ice incubation viral pre-binding step, cells were washed with cold PBS (pH = 7.4) to remove unbound viral particles, replaced with fresh media, and incubated at 37°C, 5% CO_2_. After 2h, cells were washed with high-pH PBS to remove the surface-bound virus, replaced with fresh media, and returned to 37°C, 5% CO_2_. Samples were then collected at the indicated times and processed for non-reducing SDS-PAGE. For some binding entry experiments, 20mM final concentration of NH_4_Cl were added to prevent viral degradation.

### L2-BirA membrane penetration assay

60,000 cells/well of HaCaT-GFP-BAP cells were plated in a 24-well plate. Cells were infected with 150ng L1/well of HPV16 L2-BirA virus with 5μM or 10μM peptides. At 24h post-infection, samples were processed for reducing SDS-PAGE followed by western blot to detect total and biotinylated GFP.

### Aphidicolin release experiment

50,000 cells/well of HaCaTs were seeded in 24 well plates. Cells were then infected with HPV16 at 2x10^8^ viral genome equivalents (pGL3 copies) per well under the presence of 6μM aphidicolin. At 18h post-infection, cells were washed with high pH PBS (pH=10.6-10.8) for 2.5 mins followed with two washes in regular PBS (pH=7.2-7.4) to remove all the surface-bound virus and neutralize the pH. Then, fresh culture media supplied with 6μM aphidicolin was added. After 8h incubation, the aphidicolin was released by replacing with fresh media.

### Subcellular fractionation and alkaline membrane extraction

2 million HeLa-S3 cells were seeded in a 6cm dish. The following day cells are infected with 1.25x10^5^ virions/cell at 37°C, 5% CO_2_ for 24h in the presence of 10μM PTD4 or SNX1.3 and 6μM aphidicolin to prevent viral trafficking beyond the Golgi. At the end of the infection cells are rinsed with PBS and overlaid with 680μl swelling lysis buffer (10mM HEPES pH=7.5, 1.5mM MgCl_2_ hexahydrate, 10mM KCl, 0.5mM DTT, 1x protease inhibitor cocktail (Sigma P8340), 1x PhosStop cocktail (Sigma) on ice for 30m. Cells were collected by scraping and transferred to a 2ml Dounce homogenizer on ice. Samples were homogenized with 30 full strokes of pestle B and transferred to microfuge tubes. At this time, an 80μl aliquot of the sample corresponding to the “T total” fraction was taken and combined with 20μl SDS-PAGE loading buffer and incubated at 90°C for 7m. The remaining sample was centrifuged at 10,000xg for 10 min at 4°C. Nuclear pellets were discarded and 550μl the supernatant was transferred to 5x41mm ultra clear centrifuge tubes (Beckman Coulter 344090). Samples were balanced and centrifuged at 42,000 rpm for 35m at 4°C in a SW55Ti rotor and buckets with tube adaptors (Beckman REF: 356860). The supernatant was then concentrated to 80μl by spinning through Amicon 3kDa MWCO concentrator units at 14,000xg for 15-30m at 4°C. These “S1 cytosol” fractions were then combined with 20μl SDS-PAGE loading buffer and heated to 90°C for 7m. The pellets were gently rinsed (not resuspended) with 600μl HN buffer (50mM HEPES pH=7.5, 150mM NaCl) and centrifuged again at 42,000 rpm for 35m at 4°C. The supernatant is discarded and the pellet from this step is rinsed and resuspended in 50μl alkali buffer #1 (10mM Tris-HCl pH=7.5, 150mM NaCl, 2mM MgCl2 hexahydrate, 5mM DTT) for 15m on ice, at which time 500μl alkali buffer #2 (100 mM Na_2_CO_3_ pH=11.7, freshly prepared) and samples are incubated for 30m on ice. Samples were then spun at 42,000 rpm for 35m at 4°C to pellet the membranes. Supernatants were collected and concentrated to 80μl by spinning through Amicon 3kDa MWCO concentrator units at 14,000xg for 15-30m at 4°C. These “S2 luminal and peripheral proteins” were then mixed with 20μl SDS-PAGE loading buffer and heated to 90°C for 7m. Membrane pellets were rinsed with 600μl HN buffer and spun again at 42,000 rpm for 35m at 4°C. Supernatant was carefully removed and discarded, and these final “P membrane” fractions were dissolved in 50μl SDS-PAGE loading buffer and heated to 90°C for 7m. All fractions (T, S1, S2, P) were analyzed by SDS-PAGE and western blotting for HA (L2), Vps35 (cytosolic retromer), and BAP31 (ER membrane), as described above.

### Structural modeling, sequence alignment, and STRING analysis

The SNX1/SNX5 heterodimer (PDB 8AFZ) was rendered with PyMol [64], highlighting the portion of SNX1 corresponding to the SNX1.3 peptide in red. The full structure is the ESCPE-1 membrane coat [56], which contains a portion of the cation-independent mannose phosphate receptor (CI-M6PR) in complex with the SNX1/SNX5 heterodimer. The CI-M6PR density was removed for clarity. MacVector software was used for sequence alignment of SNX1.3 and SNX2.1 peptides, using the ClustalW algorithm [65]. STRING interaction network analysis was performed using STRING version 12.0 [66] webserver: [67]. “SNX1”, “SNX2”, and “EGFR” were used as the input queries.

## Results

### SNX1.3 blocks HPV16 infection in multiple cell lines

Infection experiments were performed using HPV16 pseudovirus encapsidating a firefly luciferase reporter plasmid as described in *Materials and Methods*. SNX1.3 or control PTD4 peptides were added at the time of infection and luciferase was assayed 24h post infection. SNX1.3 treatment resulted in a dose-dependent inhibition of HPV16 infection of HaCaT, 293TT, and murine K38 keratinocytes, whereas PTD4 control peptide had no significant effects (Fig 1A, B). Calculated IC_50_ values were 3.8μM for 293TT, 2.6μM for HaCaT, and 2.9μM for K38 keratinocytes (Fig 1C). Nearly two logs of inhibition were observed at the 10μM dose in all three cell lines (Fig 1B, C). Mitosis is a requirement for HPV16 infection [68], and 10μM SNX1.3 peptide was cytotoxic to triple negative breast cancer cells [57], so it is important to determine if the peptide affects cell proliferation or viability over these timeframes. 10μM PTD4 and SNX1.3 had only minor effects on cell proliferation, extending the calculated doubling time of HaCaT cells from 24.7h hours to 25.8h and 28.2h respectively, as determined from exponential growth curve fitting of total cell counts after 72h of continuous culture in the presence of peptides (Fig 1D). The differences in cell proliferation were particularly minimal in the first 24h of growth, the timeframe used for most experiments in this work. No significant differences in total cell count were observed between the groups at any timepoint as determined by 2-way ANOVA. SNX1.3 had no evidence of cytotoxicity compared to PTD4, as measured by WST-1 assay (Fig 1E). Taken together, the data show that SNX1.3 effectively blocks HPV16 pseudovirus infection in multiple cell types, and that this inhibition is not simply due to general cytotoxicity or a block of mitosis.

### SNX1.3 peptide delays HPV16 entry

To understand SNX1.3 inhibition of HPV16 we began with binding and entry assays. For binding assays HaCaT cells were chilled to 4°C and incubated with virus for 1h in the presence of PTD4 or SNX1.3 peptides. Unbound virus was washed away with PBS and some samples were treated with a high pH PBS buffer (pH = 10.75) to remove bound virions [24]. Cells underwent additional PBS rinses and proteins were extracted for non-reducing SDS-PAGE. Bound virions were detected by western blot for the disulfide-bonded dimeric and trimeric forms of the major capsid protein L1. Figure 2A is a representative blot, with quantified data including additional biological replicates shown in Figure 2B. The data clearly shows that SNX1.3 has no significant impact on virion binding and that high pH washing removes nearly all the surface bound virus, which is important for the entry assays described below.

**Fig 2.**
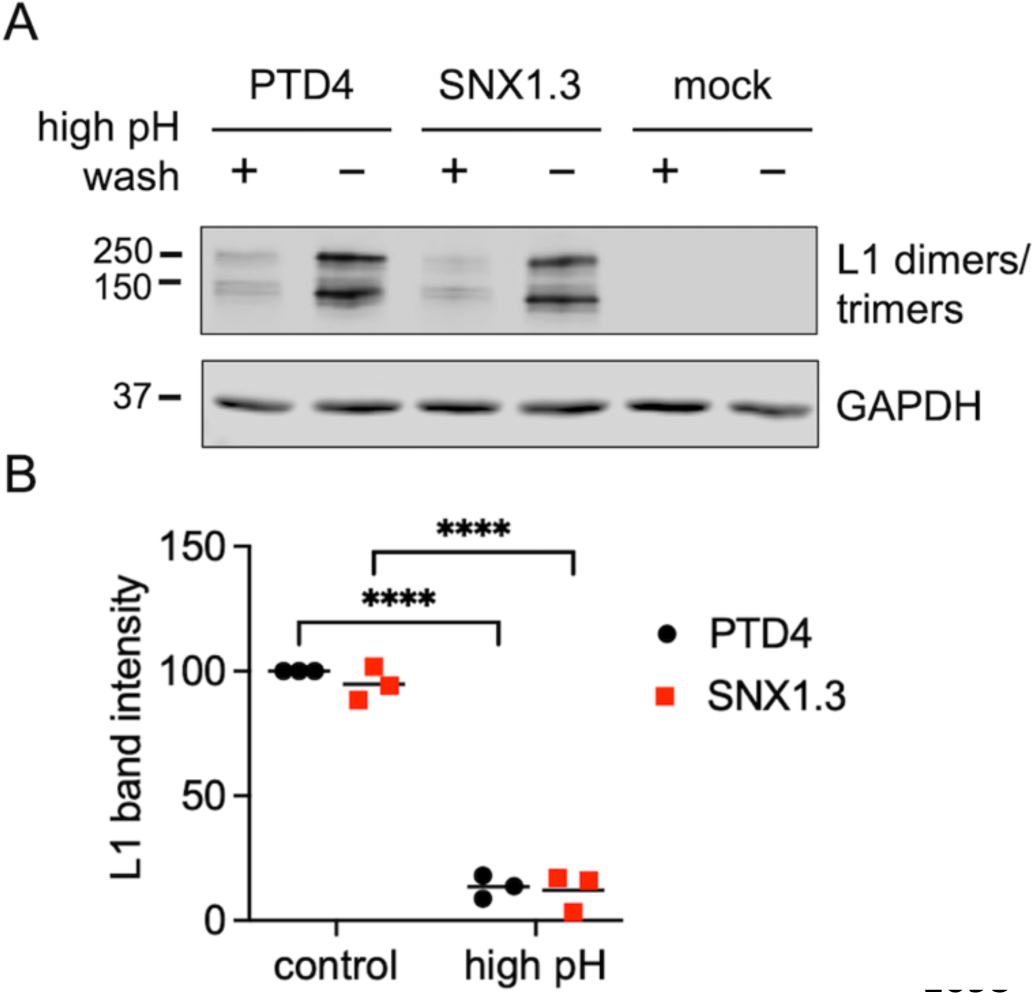
HPV16 binding assay. **(A)** HPV16 was bound to HaCaT cells ± 10uM PTD4 or SNX1.3 for 1h at 4°C. Cells were then washed with regular PBS to remove unbound virus in the media or washed with high pH PBS to remove the surface-bound virus as well. Cell surface-bound L1 dimers and trimers were detected by non-reducing SDS-PAGE and western blot, as described in *Materials and Methods*. A representative L1 blot is shown, along with GAPDH loading control. **(B)** Densitometric quantification of intact HMW L1 band intensities was performed by ImageJ. Graphs show mean relative L1 band intensities at given times, n = 3 independent biological replicates, relative to the PTD4 treated groups. Statistics were determined using 2-way ANOVA with Šídák’s multiple comparison test. ****p<0.0001.

Next, we investigated the effects of SNX1.3 on viral endocytosis. In our initial experiments, virions were prebound to HaCaT cells followed by a shift to 37°C for 3h, to allow viral entry. After the 3h incubation, extracellular virus was cleared by high pH PBS wash, and samples were either processed for non-reducing SDS-PAGE or given fresh media and returned to 37°C for another 3h before processing for SDS-PAGE. Upon return to 37°C, this “3h wave” of incoming virus will continue trafficking through the endolysosomal pathway. As the virus traffics, L1 is cleaved by acid-dependent endosomal proteases, resulting in the appearance of smaller degradation products [19,69,70]. The data show lower levels of both intact L1 and degraded L1 upon treatment with SNX1.3 compared to PTD4 (Supplemental Fig 1). Steady state levels of intracellular L1 depend on both the rates of virion uptake and the rates of virion degradation. Therefore, the observed differences in steady state L1 levels caused by SNX1.3 could either be a result of slower uptake or accelerated degradation.

To differentiate between these two possibilities, and provide a means of obtaining more quantitative data, we chose to eliminate the endo/lysosomal degradation of virions by addition of NH_4_Cl to the culture, which acts to buffer against endosomal acidification to prevent activation of endosomal proteases [19,71]. To address the kinetics of viral uptake, infections were done over a 6h time course in the presence of NH_4_Cl. A high pH wash was done at the end of each infection to measure only the intracellular virions at the indicated times. The data show potent inhibition of HPV16 entry at 2h by SNX1.3 compared to PTD4 (Fig 3A, B). The lack of degraded L1 indicate show that the NH_4_Cl treatment was effective (Fig 3A) allowing us to measure cumulative entry. Over time, the internalized virus increased steadily for both the PTD4 and SNX1.3 groups (Fig 3B). Plotting the best fit linear regression of the data revealed nearly identical slopes (Fig 3C), indicating that SNX1.3 treatment substantially blunted early entry, but over time virions enter at the same rate as the PTD4 group. The net result appears to be a delay in viral uptake.

**Fig 3.**
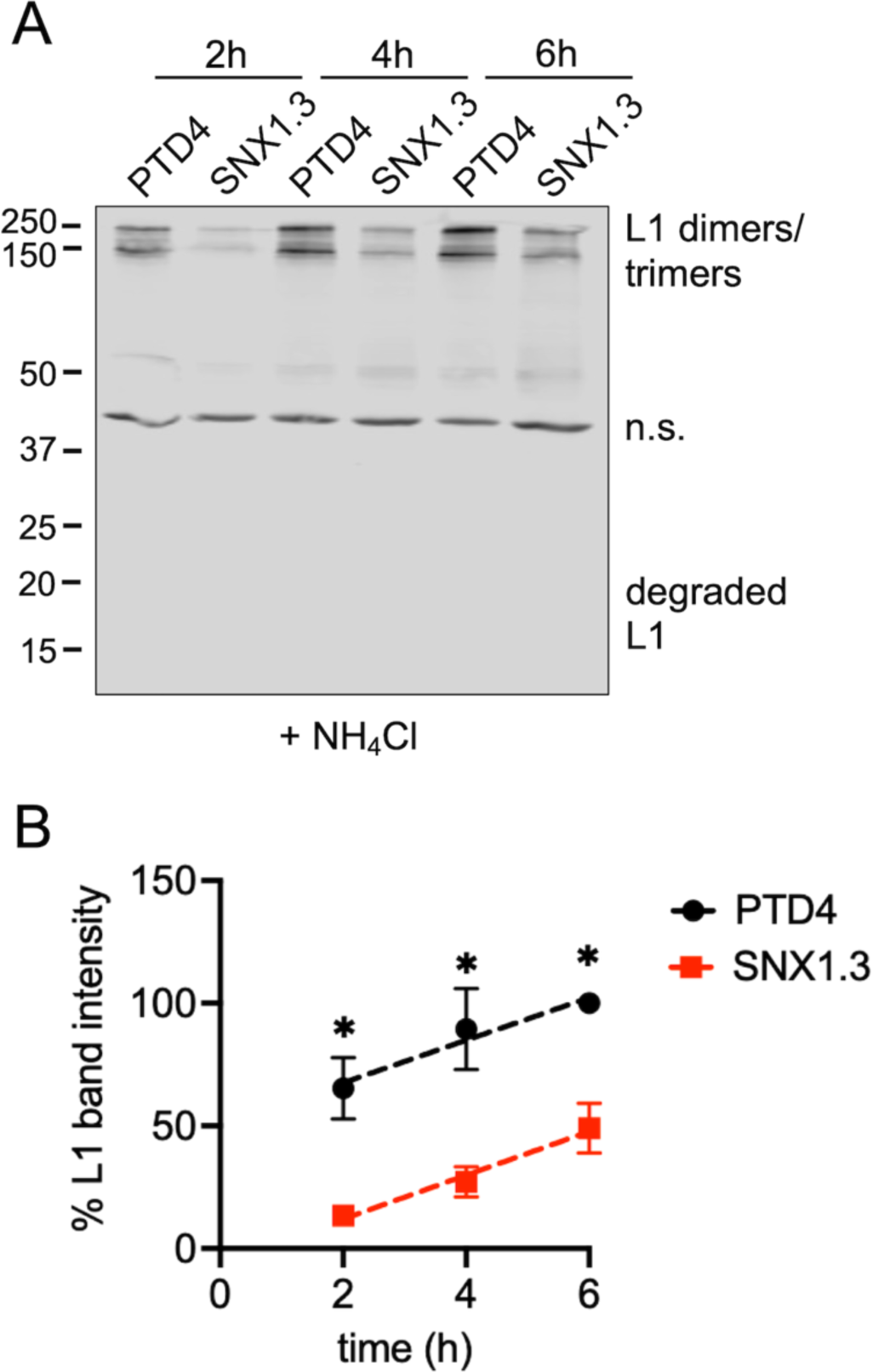
SNX1.3 delays HPV16 entry. **(A)** HPV16 was bound to HaCaT cells ± 10μM PTD4 or SNX1.3 and 20 mM NH_4_Cl for 1h at 4°C. Cells were then incubated at 37°C for 2h, 4h, or 6h at which time cell surface virus was removed by high pH PBS wash at the time of sample collection. Cell lysates were processed for non-reducing SDS-PAGE and western blot to detect intact and degraded forms of L1. Non-specific bands (n.s). is a cellular protein that cross-reacts with the L1 antibody and serves as an internal loading control. In this manner cumulative viral uptake can be visualized as more and more virions enter the cells over the timecourse and degradation is blocked by NH_4_Cl. Representative blot is shown. **(B)** Densitometric quantification of intact HMW L1 band intensities was performed by ImageJ. Graphs show mean relative L1 band intensities at given times (±SEM, n = 3 independent biological repeats) relative to the PTD4 treated group at 6h time point. Simple linear regression was used to plot the lines. Statistics were determined using multiple unpaired t-tests. *p<0.05.

To confirm this phenotype, we performed immunofluorescence experiments, directly visualizing intracellular virions by confocal microscopy. Cells were infected in the presence of NH_4_Cl and either PTD4 or SNX1.3 for 2h, 4h, or 18h prior to removal of surface virus with high pH PBS, additional washing, fixation, and immunofluorescent staining for the capsid and the endolysosomal markers CD63 or LAMP1. The representative micrographs again show relatively poor uptake of virions in the presence of SNX1.3 at early time points, but eventually the intracellular levels of capsid within SNX1.3 treated cells become equivalent to those of the PTD4 treated group (Supplemental Fig 2A). The micrographs also show that intracellular viruses are colocalizing with endolysosomal markers in both groups, although the colocalization with LAMP1 at 18h appears to be altered upon SNX1.3 treatment, perhaps suggestive of altered trafficking. Quantification and normalization of the intracellular capsid signal to cell number confirm the initial delay of viral uptake in the SNX1.3 treated groups (Supplemental Fig 2B,C,D). Over time, the average intracellular levels of HPV16 virion in the SNX1.3 treated group reach and even slightly surpass those of the PTD4 treated group (Supplemental Fig 2D). The fact that SNX1.3 causes such a profound inhibition of HPV16 infection suggests that SNX1.3 must be causing a secondary block to subcellular viral trafficking, downstream of entry. Moving forward, we decided to focus on this aspect of SNX1.3 activity.

### Late mitotic trafficking and limiting membrane penetration is blocked by SNX1.3

Successful HPV16 infection requires the delivery of the viral genome to the host cell nucleus, a process dependent on mitosis [40,68,72]. During late mitotic trafficking, vesicles containing the L2/vDNA complex traffic along microtubules, tethering on host mitotic chromosomes via the central chromatin binding domain (residues 188-334) within L2 [41]. The chromosome-bound vesicular L2/vDNA ends up in daughter cell nuclei and the limiting vesicular membrane is quickly lost during this M/G1 transition [39]. Loss of this vesicular membrane is important for the vDNA to access the cellular transcriptional and replicative machinery necessary for successful infection. In prior work we developed an assay for this late limiting membrane penetration, which we termed “membrane translocation” at the time [40]. The assay is based on the compartmentalization of the biotin-protein ligase BirA [73] away from its ectopically expressed cytosolic substrate, GFP fused to a short biotin acceptor peptide [74](GFP-BAP). Infection of GFP-BAP expressing HaCaT cells with HPV16 virions packaging an L2-BirA fusion results in biotinylation of GFP-BAP only if L2-BirA can fully breach the limiting membrane barrier during late mitotic trafficking. We first confirmed that SNX1.3 was inhibitory to L2-BirA virion infection of HaCaT-GFP-BAP cells (Fig 4A). Next, we assayed the penetration of limiting membranes by detection of biotinylated GFP-BAP using SDS-PAGE and western blot with IR800-conjugated fluorescent neutravidin. The data show a strong GFP-biotin signal in PTD4 treated cells, but a strong reduction of GFP-biotin in the presence of 5 μM and 10 μM SNX1.3 (Fig. 4B, C). Densitometry shows a 75% and 95% decrease of L2-BirA membrane penetration with 5 μM and 10 μM SNX1.3 (Fig. 4C), suggesting that the very late stages of nuclear trafficking and penetration of limiting membranes are blocked by SNX1.3.

**Fig 4.**
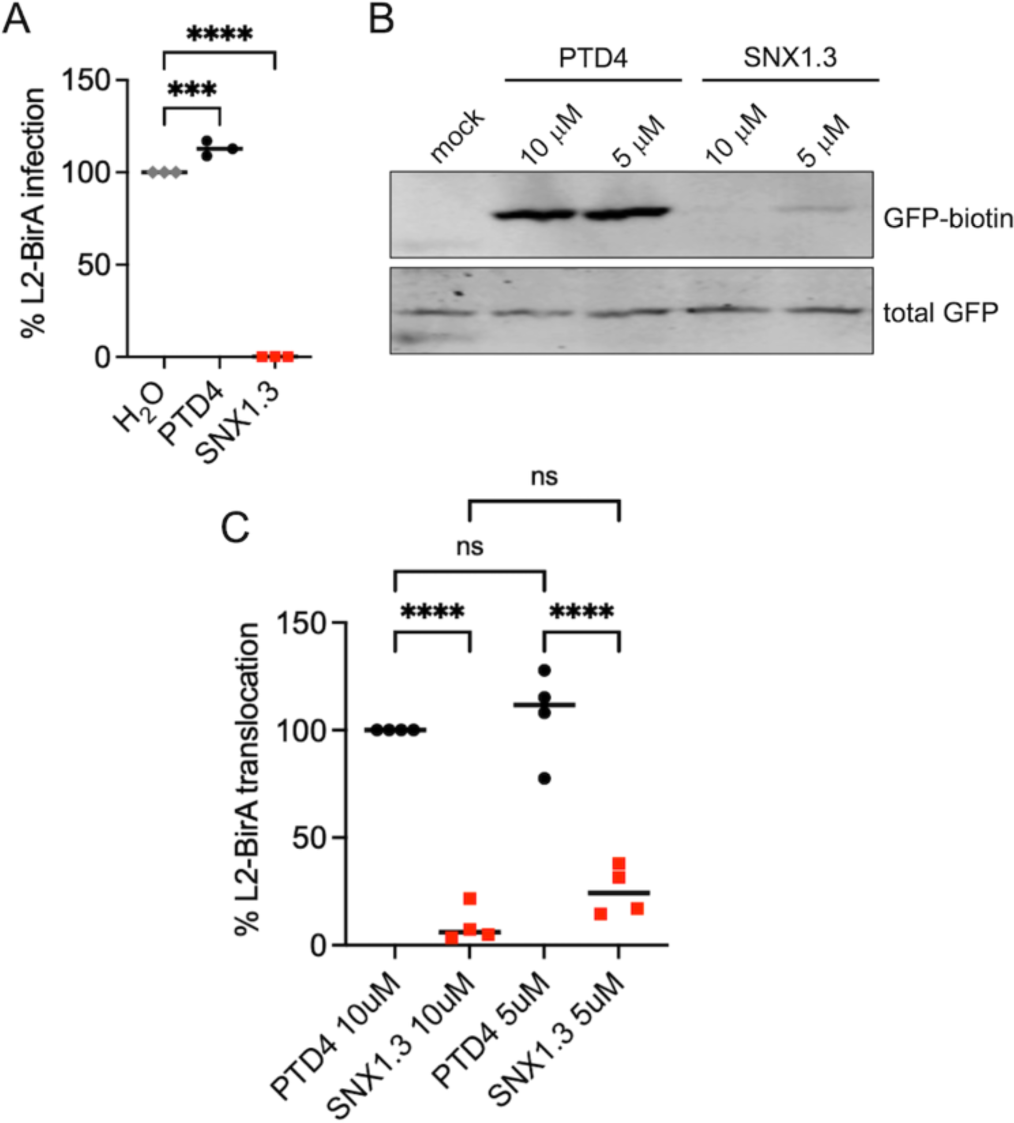
Late membrane penetration is blocked by SNX1.3. **(A)** Infectivity of luciferase-expressing HPV16 packaged with L2-BirA with 10μM PTD4, SNX1.3, or H2O (solvent vehicle control) in the HaCaT-GFP-BAP cell line. **(B)** L2-BirA membrane translocation assay. HaCaT-GFP-BAP cells were infected with HPV16 bearing L2-BirA with 10μM PTD4 or SNX1.3 peptides. Cells were processed for SDS-PAGE and western blotting at 24h post-infection, blotting for biotin-GFP and total GFP. Biotinylation of GFP can only occur if L2-BirA penetrates the limiting vesicular membrane during late mitotic trafficking of virions **(C)** Quantification of the membrane translocation assay. Band intensities were measured by densitometry using ImageJ. The 10μM PTD4 treated cells were set to 100%, The graph represents the mean percent biotin-GFP normalized with total GFP ±SEM, n = 4 independent biological replicates. Statistics were determined using 1-way ANOVA with Šídák’s multiple comparison test. ****p<0.0001.

### SNX1.3 blocks retrograde transport of HPV16 but does not affect post-Golgi trafficking

As late trafficking and limiting membrane penetration are contingent on successful retrograde transport of virions from endosomes to the Golgi [37,75], we next investigated the effects of SNX1.3 on retrograde trafficking. For these experiments we used virions packaged with HA-tagged L2 (L2-HA) to detect the presence of L2 in the Golgi by immunofluorescence and confocal microscopy. We performed infections in the presence of aphidicolin, a Polα/Pol8/Polχ inhibitor that reversibly blocks cells in early S phase [76]. Under these conditions HPV16 virions will retrograde traffic and accumulate in the Golgi as the cells never reach G2/M, when Golgi egress and late nuclear trafficking occur [77]. Cells were infected in the presence of aphidicolin and either PTD4 or SNX1.3 peptide for 24 hours prior to removal of surface virions with high pH PBS, washing, fixation, and immunostaining for L2-HA, p230 Golgi marker, and DAPI. The long 24h infection ensures that the SNX1.3-treated group has time to overcome the early delay in endocytic uptake, as can be observed from the levels of L2-HA staining (Fig 5A). The data clearly show strong accumulation of L2-HA in the p230-positive trans-Golgi network in the presence of PTD4. In contrast, SNX1.3 treatment abolished the overlap between L2-HA and p230, showing a clear defect in retrograde trafficking (Fig 5A). Combined quantitative data of the Manders’ colocalization coefficients from three independent experiments show a statistically significant impairment of L2-HA/p230 colocalization by the SNX1.3 peptide (Fig 5B).

**Fig 5.**
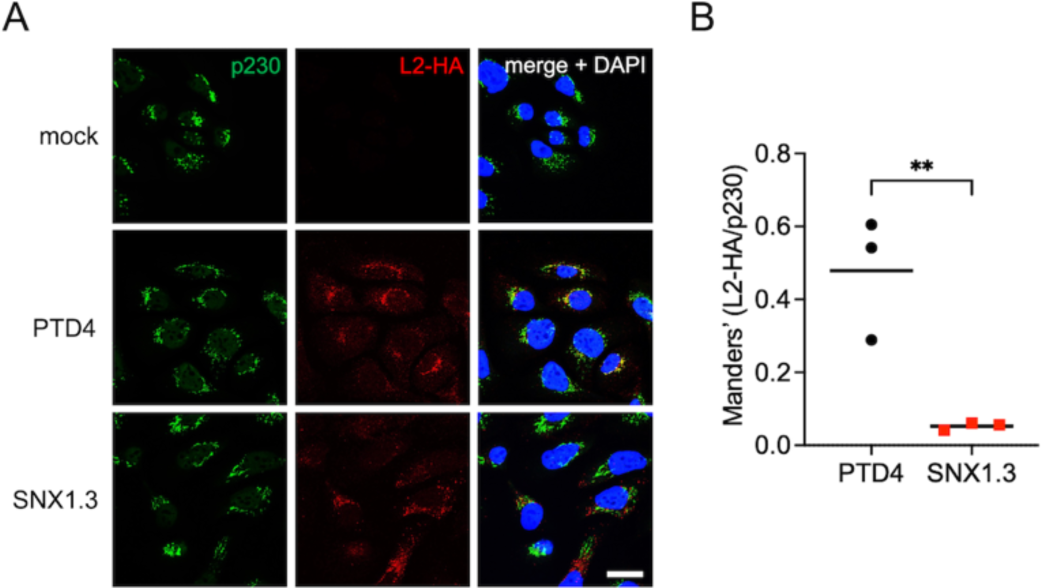
SNX1.3 blocks HPV16 retrograde trafficking to the Golgi. **(A)** HaCaT cells were infected at 37°C with HPV16 bearing L2-HA tag for 24h together with 6 μM aphidicolin and either 10μM PTD4 or SNX1.3 peptide at 37°C for 24hrs followed by a high pH wash to remove the surface-bound virus before fixation and processing for IF staining as described in *Materials and Methods*. Cells were stained with mouse anti-p230 with AlexaFluor-488 conjugated anti-mouse secondary (green), and rat anti-HA with AlexaFluor-555 conjugated anti-rat secondary (red). The cell nucleus was stained with DAPI (blue). Scale bars, 20 μm (**B)** Colocalization analysis using the JACoP plugin of ImageJ. Manders’ overlap coefficients were measured between L2-HA:p230 for multiple Z-stacks, each containing multiple cells/field, from 3 independent experiments. The mean Manders’ coefficient of each biological replicate ±SEM are shown in black bars, dots are the mean Manders’ coefficient for each biological replicate, each determined from numerous technical replicate measurements. Statistics were determined using unpaired t-tests. **p<0.01.

We further tested whether SNX1.3 is still inhibitory after the virus has already reached the *trans*-Golgi network. Aphidicolin is a reversible cell cycle inhibitor, once it is removed by washout, viruses that are trapped in the *trans*-Golgi compartments will resume their post-Golgi nuclear trafficking pathway once infected cells cycle into G2M [40,78]. The experimental setup is shown in the Fig 6A. HaCaT cells were infected with HPV16 in the presence of 6μM aphidicolin for 18h prior to high pH PBS removal of surface bound virus. Since viral uptake is not synchronized, cells were further incubated in media with 6μM aphidicolin for another 8h to allow the intracellular virions additional time to reach *trans*-Golgi network. At 26h post infecton, cells were released from aphidicolin by washing and replacing with media containing either PTD4 or SNX1.3 peptide for an additional 24h prior to luciferase assay. We did not observe any significant inhibition of infection by SNX1.3 in this assay (Fig. 6B). These data show that HPV16 fails to undergo efficient retrograde trafficking in the presence of SNX1.3, but once the virus has reached the Golgi, SNX1.3 treatment is no longer inhibitory.

**Fig 6.**
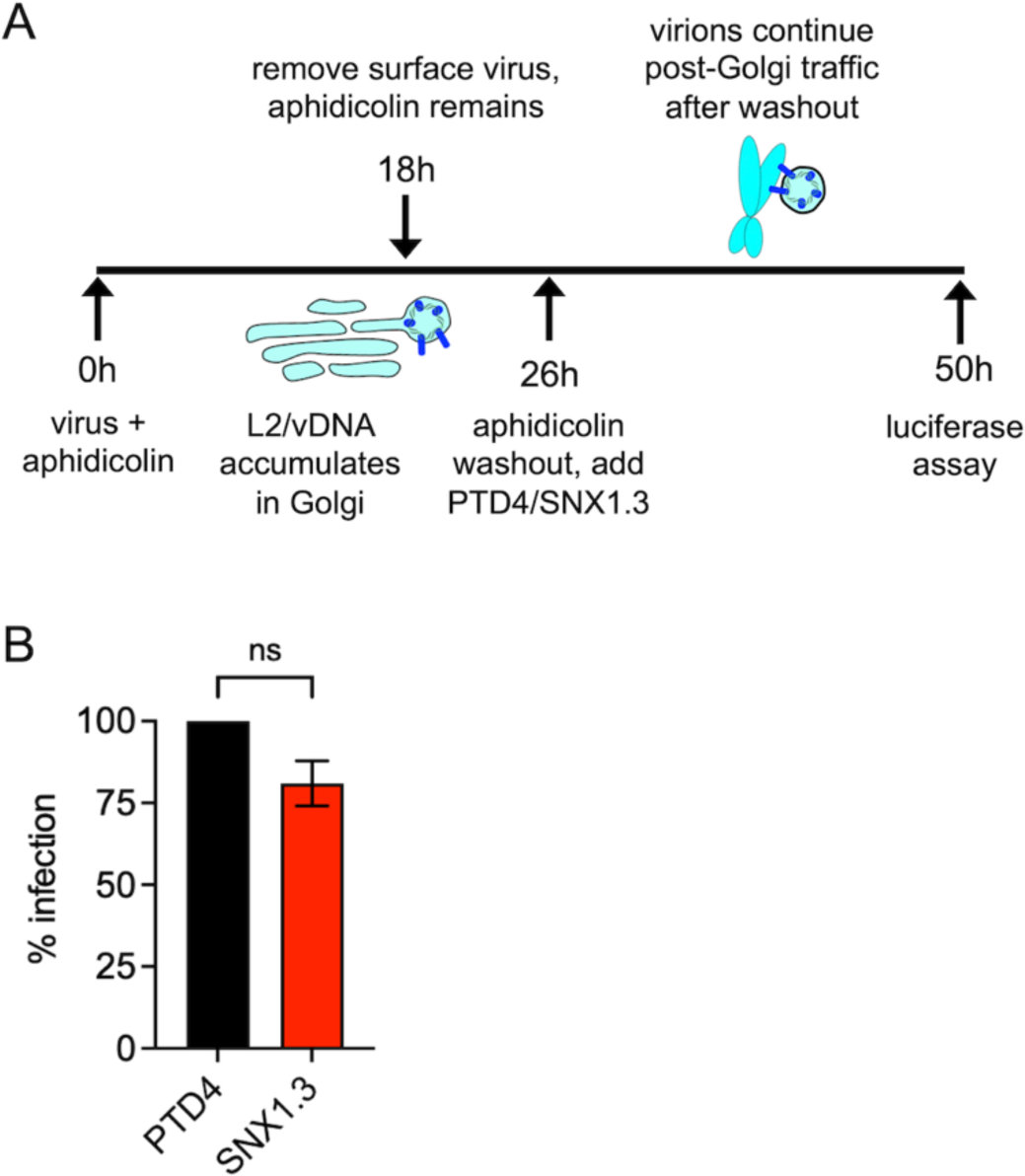
SNX1.3 has no effect on post-Golgi trafficking. **(A)** Schematic of aphidicolin release infection experiment. HaCaTs were infected with HPV16 at 37°C in the presence of 6μM aphidicolin for 18h followed with high pH PBS (pH=10.6-10.8) wash to remove all the surface-bound virus. Fresh culture media supplemented with 6μM aphidicolin was then added for another 8h incubation at 37°C. This allows for intracellular viruses to retrograde traffic to the Golgi. Aphidicolin was then released by replacing media with containing either 10 μM PTD4 or SNX1.3 peptide for another 24h at 37°C to allow nuclear trafficking of virus and the expression of luciferase. **(B)** Infectivity of luciferase-expressing HPV16 in the aphidicolin release experiment, normalized to PTD4 treatment. Mean infectivity ±SEM are plotted from n=3 biological replicates. Statistics were determined with a two-sample unpaired t-test.

### SNX1.3 inhibits L2 membrane spanning

The minor capsid protein L2 is required for the retrograde transport of the vDNA. After endocytic uptake, L2 is inserted into the local endosomal membrane to adopt a type-I transmembrane protein topology [33], spanning via its single N-terminal transmembrane domain [32]. Precise mechanisms of L2 spanning remain obscure, but the process is dependent on numerous cellular factors including furin, p120 catenin, γ-secretase, and endosomal acidification, in addition to a positively charged cell penetrating peptide near the C-terminus of L2 [30,31,79]. Membrane spanning in this manner allows ∼400 C-terminal residues downstream of the transmembrane domain to protrude into the cytosol to recruit retromer and other sorting factors necessary for efficient retrograde trafficking to the Golgi [34,35,37]. We therefore wanted to determine if the block in retrograde trafficking by SNX1.3 was due to a defect in membrane spanning.

To measure spanning we adopted an alkali membrane extraction protocol recently described by the Tsai Lab [31]. This stringent subcellular fractionation protocol relies on sequential high-speed centrifugations and a harsh alkali wash step to isolate only *bona fide* integral transmembrane proteins in the final membrane pellet. HeLa-S3 cells were infected in the presence of either PTD4 or SNX1.3 for 24h. Aphidicolin was included in the media to prevent virion traffic beyond the Golgi in this long infection. Cells were gently lysed and fractionated as described in *materials and methods*. We collected four fractions: T; total cell fraction, S1; cytosolic protein, S2; luminal organelle and peripheral membrane proteins, and P; integral membrane proteins. Samples were analyzed by SDS-PAGE and immunoblotting for L2-HA, the cytosolic marker Vps35, and the ER transmembrane protein BAP31. Results show a clear reduction in L2 membrane spanning in the presence of SNX1.3 (Fig 7A). Results from three independent experiments were quantified by densitometry, normalizing the membrane-spanning L2 signal in the P fractions to both the BAP31 membrane marker and the total L2 signal in the T fraction, revealing a strong and statistically significant defect in membrane spanning upon addition of SNX1.3 peptide (Fig 7B). These data suggest that SNX1.3 blocks retrograde trafficking of virus by somehow preventing the membrane spanning ability of L2.

**Fig 7.**
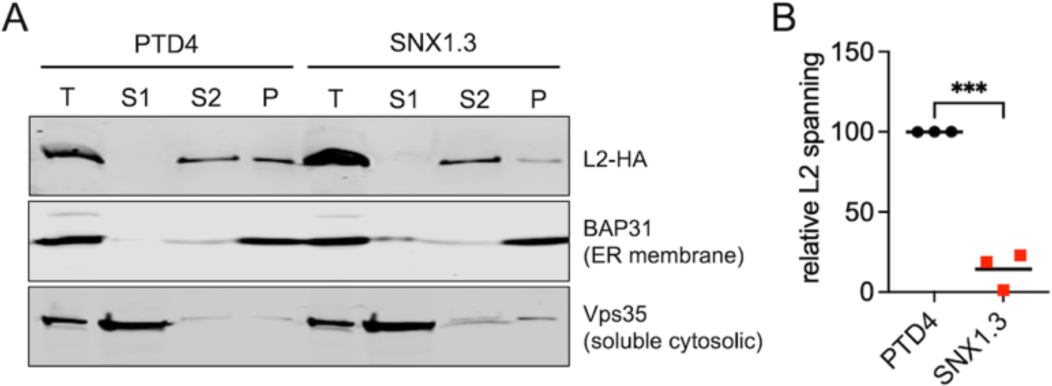
SNX1.3 blocks L2 membrane spanning. **(A)** Subcellular fractionation and alkali membrane extraction blot of membrane spanning L2. HeLa-S3 cells were infected with HPV16 bearing L2-HA in the presence of 10μM PTD4 or SNX1.3 peptide and 6μM aphidicolin as described in *Materials and Methods*. After 24h cells were processed for fractionation and alkaline membrane extraction. Total (T), S1 (supernatant 1; cytosolic proteins), S2 (supernatant 2; organelle lumenal and peripheral membrane proteins), and P (final pellet; integral membrane proteins) were collected as described and analyzed by SDS-PAGE and western blotting for BAP31 (integral ER membrane protein), Vps35 (soluble cytosolic protein), and L2-HA. Representative blot is shown. **(B)** Densitometric quantification of L2 membrane spanning. L2 densities within the membrane and total fractions were first normalized to their corresponding BAP31 densities to account for sample loss during fractionation and potential differences in gel loading. Then the membrane L2 density was divided by the total L2 density, with the PTD4 membrane L2 values being adjusted to 100. Mean relative L2 spanning values ±SEM are plotted from n=3 biological replicates. Statistics were determined with a two-tailed unpaired t-test. ***p<.005.

## Discussion

Previously, the SNX1.3 peptide was found to inhibit EGFR retrograde nuclear trafficking in TNBC cells, causing cell death [57]. Here, we investigate the inhibition of HPV16 pseudovirus infection by the SNX1.3 peptide in human HaCaT keratinocytes, 293TT cells, and mouse K38 keratinocytes. We found the peptide did not impair cell viability or proliferation of these cells, in contrast to the nuclear EGFR-dependent TNBC. SNX1.3 impaired HPV16 infection with low micromolar IC_50_ values, causing a delayed uptake phenotype in HaCaT cells. Given time, HPV16 virions eventually overcome this entry delay but SNX1.3 peptide treatment imposed a strong block to late nuclear trafficking and retrograde Golgi trafficking of L2/vDNA. Once virus reached the Golgi, infection was resistant to SNX1.3 treatment, demonstrating that the block in trafficking was pre-Golgi. Alkaline membrane fractionation experiments revealed that the block in retrograde trafficking by SNX1.3 was due to a defect in L2 membrane spanning, a necessary step for viral retrograde trafficking. We have not yet tested whether the SNX1.3 peptide is inhibitory to diverse HPV types, nor have we tested if the SNX1.3 peptide directly binds to L1 or L2 proteins of the HPV capsid.

Clearly more work is needed to determine if the delayed entry phenotype is due to EGFR-dependent mechanisms but SNX1 overexpression can alter EGFR signaling, endocytosis, retrograde trafficking, recycling, and degradation [51,80–82]. The effects of the SNX1.3 peptide on endogenous SNX1 and SNX-BAR function remain to be determined but it could be envisioned that the peptide may act as a dominant negative form of SNX1 in some contexts and/or excess peptide could mimic SNX1 overexpression in other contexts. These details remain to be determined experimentally but it is worth noting that the peptide is derived from a region of the SNX1 BAR homo-/heterodimerization interface and may outcompete these natural interactions (Fig 1A), and many networks and interactions with cellular partners could be perturbed by the introduction of such a peptide into the system (Supplemental Fig 3). The patterns of endolysosomal markers CD63 and LAMP1 appeared to be slightly altered by SNX1.3 treatment (supplemental Fig 2), suggesting that SNX1.3 treatment may result in a global (but subtle) perturbation of vesicular trafficking and/or altered localization of cellular proteins which may affect HPV16 trafficking.

It is also notable that this region of SNX1 has a high degree of amino acid identity with the corresponding region of SNX2 (Fig 8A) and it is reasonable to posit that SNX1.3 may interfere with SNX2-partner interactions via this conserved region. Indeed, we found the analogous SNX2-derived peptide SNX2.1 displayed strong inhibition of HPV16 infection (Fig 8B) although further work is needed to validate the same delayed entry phenotype and retrograde trafficking blocks. SNX2 was recently described to contribute to HPV16 endocytosis via a macropinocytosis-like EGFR/Abl2/WASH/actin-dependent mechanism [83]. In that work SNX2 was found to be present at HPV16 entry sites and depletion of SNX2 by siRNA was shown to inhibit viral uptake [83].

**Fig 8.**
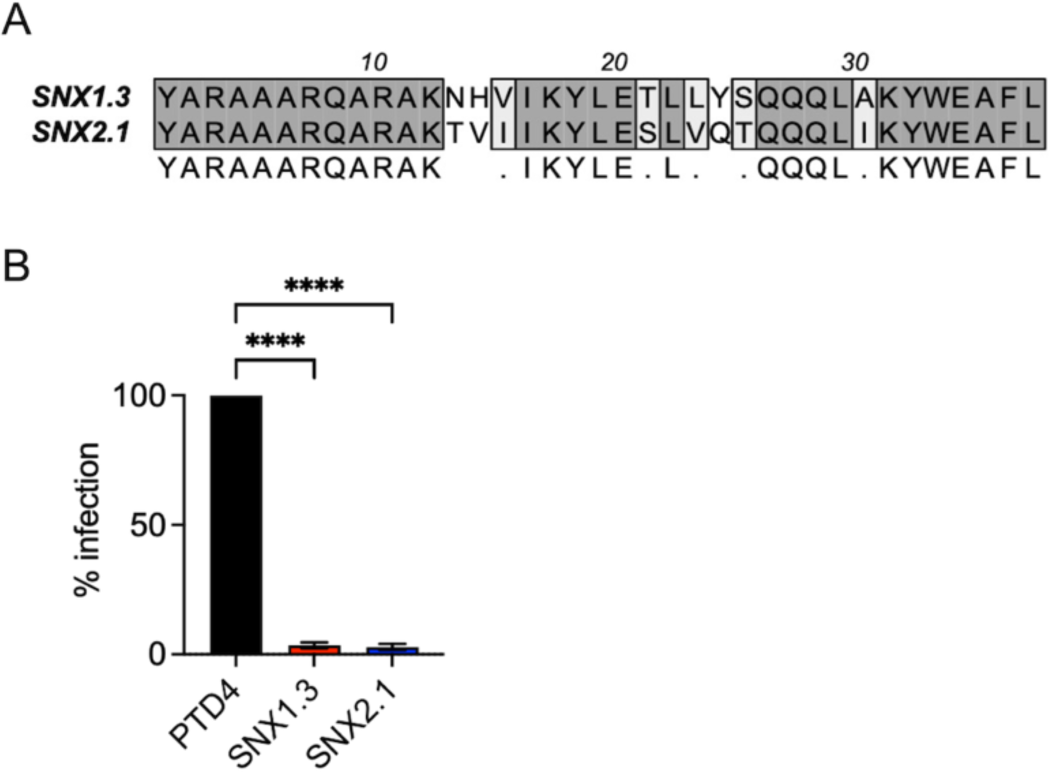
Inhibition of HPV16 infection by SNX2.1 peptide. **(A)** Sequence alignment of the SNX1.3 and SNX2.1 peptides. **(B)** Infectivity of HPV16 in HaCaT cells in the presence of 10 μM PTD4, SNX1.3, or SNX2.1 peptides. Graphs show mean infectivity ± SEM. N=3 biological replicates. Statistics were determined by 1-way ANOVA with Dunnett’s multiple comparison, ****p<0.0001.

The observed entry defect upon SNX1.3 treatment was merely a delay, with virions eventually overcoming this block. The strong block imposed by SNX1.3 was instead due to inhibition of retrograde trafficking. After entry, L2 adopts a membrane spanning conformation to gain access to cytosolic retromer and other sorting factors needed for efficient transport to the Golgi [28]. As SNX1/2/5/6 heterodimers can partner with retromer for retrograde transport of transmembrane cargo [47,52,84,85] we expected to observe normal L2 membrane spanning with a defect in transport upon SNX1.3 treatment. Rather, we observed a strong defect in L2 membrane spanning, suggesting involvement of SNX-BAR proteins or cellular partners in this process. The retromer complex was recently shown to play a role in stable membrane insertion of L2—early membrane spanning was stabilized by the binding of cytosolic retromer to its binding sites near the C-terminus of L2 [86]. Further work is needed to determine if SNX1 and related SNX-BAR proteins augment this retromer-dependent function to support L2 membrane insertion.

Alternatively, it may be that membrane tubulation and curvature are involved. HPV16 and diverse HPV types induces endosomal tubulation during entry, in an EGFR and VAP-dependent manner [87,88]. SNX2 is a known partner of VAPB and SNX2 partners with SNX1 to promote endosomal tubulation and sorting of cargo into autophagosomes upon nutrient starvation [89]. Tubulation of endosomes may generate positive membrane curvature on the lumenal face of endosomes, to aid in the membrane penetration of the C-terminal L2 cell penetrating peptide [79] and/or transmembrane domain [32]. Membrane curvature has been demonstrated to augment the activity of octaarginine cell penetrating peptide [90] and local membrane environment may greatly affect the ability of proteins like L2 to insert and span across membranes [91]. Further work is necessary to delineate the exact mechanisms behind the SNX1.3-imposed blocks we observe in this study, but this work lays the groundwork for some interesting testable hypotheses regarding potential roles of SNX-BAR proteins and their cellular partners in HPV16 infection.

## Acknowledgements.

This work was supported by grants 1R01GM136853-01 and 1R35GM152143-01 from the National Institute of General Medical Sciences. We are grateful to Dr. Joyce Schroeder and Dr. Ben Atwell for the PTD4, SNX1.3, and SNX2.1 peptides and fruitful scientific discussions. We thank Dr. Michelle Ozbun for the anti-HPV16 polyclonal antibody. We thank Dr. Koenraad van Doorslaer and members of the Van Doorslaer Lab and the Campos Lab for valuable scientific discussions.

## Author Contributions

SKC and SL designed and conceived the experiments. SL, ZLW, MAC, AJ, and SKC performed the experiments, analyzed the data, and prepared figures. SKC and SL wrote the paper.

---

**Supplemental Fig 1.**
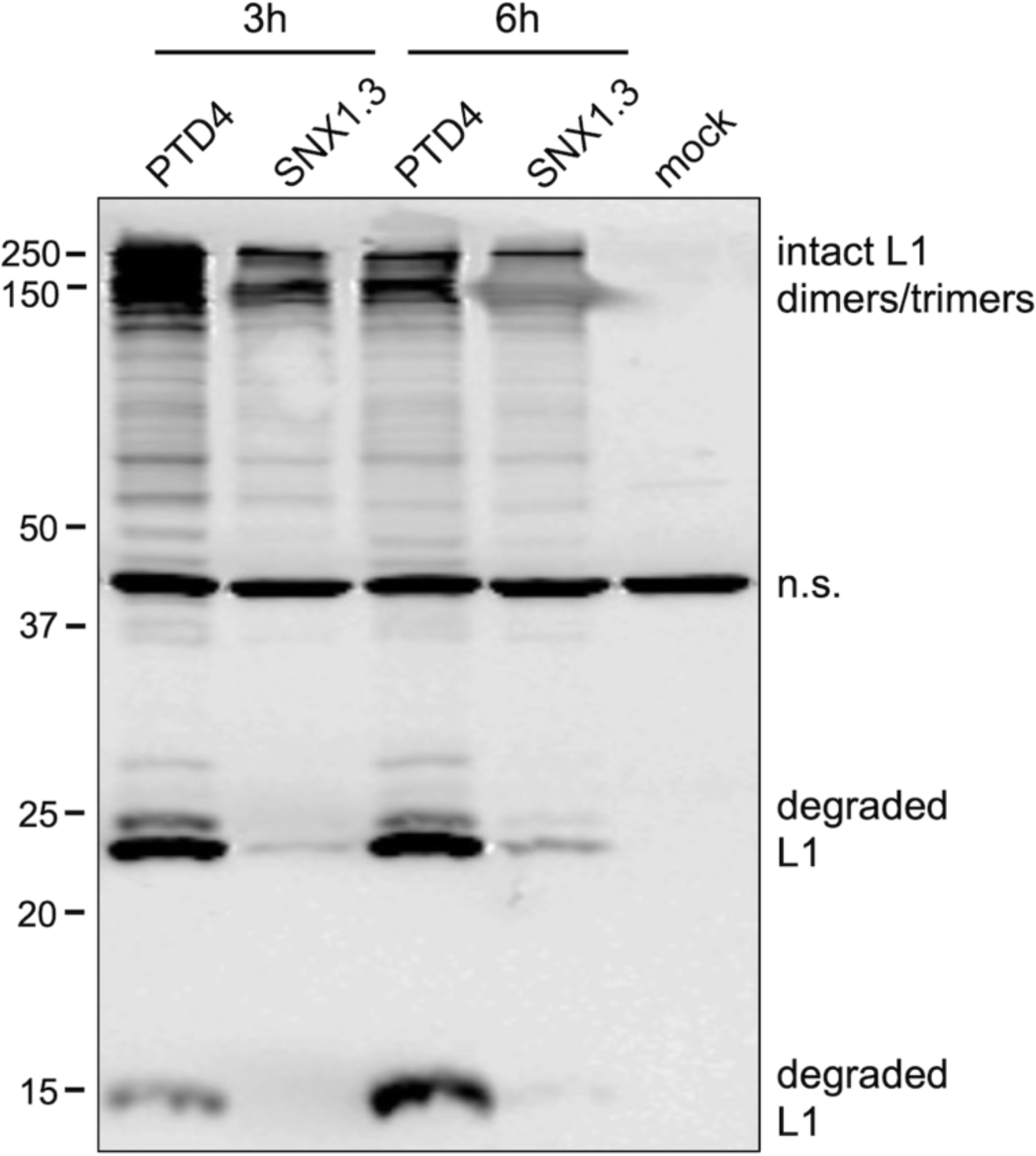
HPV16 entry assay, 3h wave. HPV16 was bound to HaCaT cells ± 10μM PTD4 or SNX1.3 for 1h at 4°C. Cells were then incubated for at 37°C for 3h to allow viral entry. At this time cell surface virus was removed by high pH PBS wash in all the groups. Media containing ± 10μM PTD4 or SNX1.3 peptides was replaced, and cells were incubated at 37°C for another 3hrs (6h time group) to allow the existing intracellular virus to traffic through the endocytic pathway. Samples were processed as described above and collected at 3h or 6h post-entry. Cell lysates were processed for non-reducing SDS-PAGE and western blot to detect intact and degraded forms of L1. Non-specific bands (n.s.) is a cellular protein that cross-reacts with the L1 antibody, and serves as an internal loading control.

**Supplemental Fig 2.**
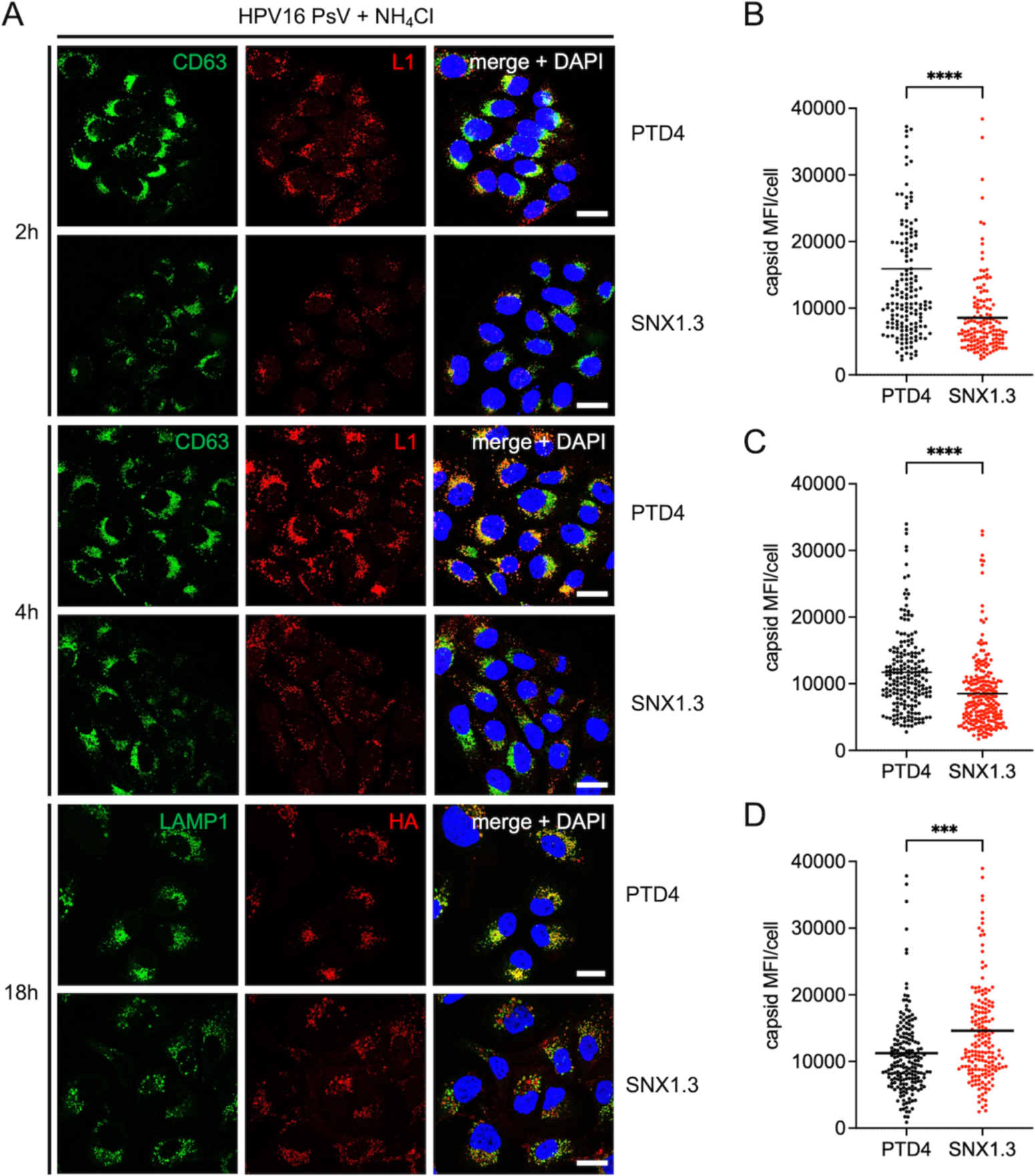
Immunofluorescence entry assay. **(A)** HaCaT cells plated on glass coverslips were infected with HPV16 bearing L2-3xFTHA ± 10μM PTD4 or SNX1.3 and 20 mM NH_4_Cl for 2h, 4h, or 18h followed by high pH PBS wash to remove the surface-bound virus. Cells were then processed for IF and capsid, endolysosomal markers, and nuclei were stained and mean fluorescence intensity of intracellular capsid signal was quantified as described in *Materials and Methods*. **(B, C, D)** Mean and all individual data points are plotted from n=3 biological replicates. Statistics were determined using multiple unpaired t-tests. ***p<0.005, ****p<0.0001. Scale bars, 20 μm.

**Supplemental Fig 3.**
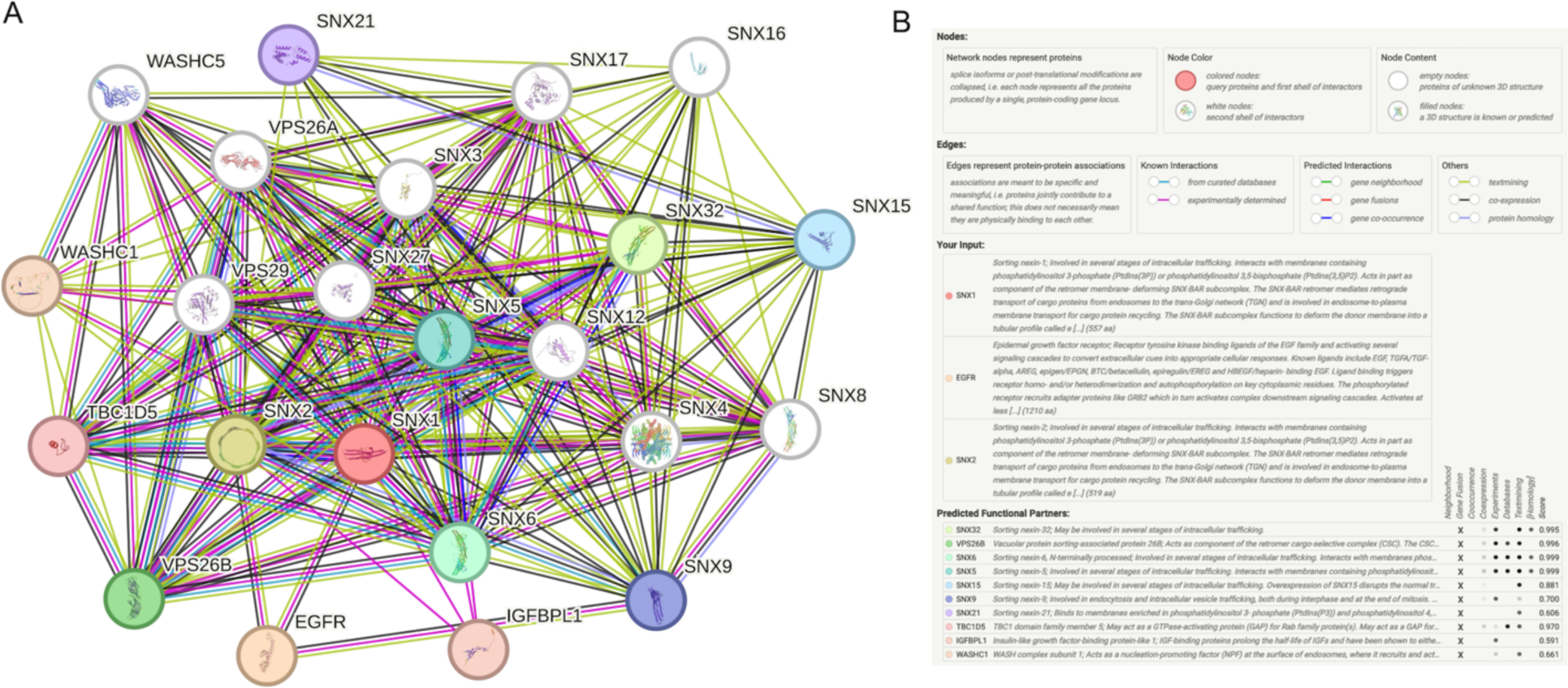
STRING network analysis of SNX1/SNX2/EGFR. **(A)** STRING version 12.0 was used to analyze the interaction network of SNX1, SNX2, and EGFR. Nodes include other SNX-BAR family members, other SNX proteins, retromer components, WASH components, and the Rab GTPase activating protein TBC1D5, many of which are cellular proteins known to be important for HPV16 trafficking and infection. **(B)** Screenshot of the network analysis legend, explaining the nodes, edges, input, and predicted partners.

## References

1. HPV Facts & Brochures | CDC. 2022,

2. Van Doorslaer K, Li Z, Xirasagar S, Maes P, Kaminsky D, Liou D, Sun Q, Kaur R, Huyen Y, McBride AA: The Papillomavirus Episteme: a major update to the papillomavirus sequence database. Nucleic Acids Res 2017, 45:D499–D506.

3. HPV and Cancer - NCI. 2019,

4. Forman D, de Martel C, Lacey CJ, Soerjomataram I, Lortet-Tieulent J, Bruni L, Vignat J, Ferlay J, Bray F, Plummer M, et al.: Global burden of human papillomavirus and related diseases. Vaccine 2012, 30 Suppl 5:F12–23.

5. Schiffman M, Doorbar J, Wentzensen N, de Sanjosé S, Fakhry C, Monk BJ, Stanley MA, Franceschi S: Carcinogenic human papillomavirus infection. Nat Rev Dis Primers 2016, 2:16086.

6. Lechner M, Liu J, Masterson L, Fenton TR: HPV-associated oropharyngeal cancer: epidemiology, molecular biology and clinical management. Nat Rev Clin Oncol 2022, 19:306–327.

7. Brotherton JML, Ogilvie GS: Current status of human papillomavirus vaccination. Curr Opin Oncol 2015, 27:399–404.

8. Zhai L, Tumban E: Gardasil-9: A global survey of projected efficacy. Antiviral Res 2016, 130:101–109.

9. Buck CB, Day PM, Trus BL: The papillomavirus major capsid protein L1. Virology 2013, 445:169–174.

10. Buck CB, Cheng N, Thompson CD, Lowy DR, Steven AC, Schiller JT, Trus BL: Arrangement of L2 within the papillomavirus capsid. J Virol 2008, 82:5190– 5197.

11. Wang JW, Roden RBS: L2, the minor capsid protein of papillomavirus. Virology 2013, 445:175–186.

12. Campos SK: Subcellular Trafficking of the Papillomavirus Genome during Initial Infection: The Remarkable Abilities of Minor Capsid Protein L2. Viruses 2017, 9:E370.

13. Doorbar J, Quint W, Banks L, Bravo IG, Stoler M, Broker TR, Stanley MA: The biology and life-cycle of human papillomaviruses. Vaccine 2012, 30 Suppl 5:F55–70.

14. Kines RC, Thompson CD, Lowy DR, Schiller JT, Day PM: The initial steps leading to papillomavirus infection occur on the basement membrane prior to cell surface binding. Proc Natl Acad Sci U S A 2009, 106:20458–20463.

15. Shafti-Keramat S, Handisurya A, Kriehuber E, Meneguzzi G, Slupetzky K, Kirnbauer R: Different heparan sulfate proteoglycans serve as cellular receptors for human papillomaviruses. J Virol 2003, 77:13125–13135.

16. Culp TD, Budgeon LR, Marinkovich MP, Meneguzzi G, Christensen ND: Keratinocyte-secreted laminin 5 can function as a transient receptor for human papillomaviruses by binding virions and transferring them to adjacent cells. J Virol 2006, 80:8940–8950.

17. Brendle SA, Christensen ND: HPV Binding Assay to Laminin-332/Integrin α6β4 on Human Keratinocytes. In Cervical Cancer: Methods and Protocols. Edited by Keppler D, Lin AW. Springer; 2015:53–66.

18. Becker M, Greune L, Schmidt MA, Schelhaas M: Extracellular Conformational Changes in the Capsid of Human Papillomaviruses Contribute to Asynchronous Uptake into Host Cells. J Virol 2018, 92.

19. Cerqueira C, Samperio Ventayol P, Vogeley C, Schelhaas M: Kallikrein-8 Proteolytically Processes Human Papillomaviruses in the Extracellular Space To Facilitate Entry into Host Cells. J Virol 2015, 89:7038–7052.

20. Richards RM, Lowy DR, Schiller JT, Day PM: Cleavage of the papillomavirus minor capsid protein, L2, at a furin consensus site is necessary for infection. Proc Natl Acad Sci U S A 2006, 103:1522–1527.

21. Mikuličić S, Florin L: The endocytic trafficking pathway of oncogenic papillomaviruses. Papillomavirus Res 2019, 7:135–137.

22. Mikuličić S, Strunk J, Florin L: HPV16 Entry into Epithelial Cells: Running a Gauntlet. Viruses 2021, 13:2460.

23. Ozbun MA: Extracellular events impacting human papillomavirus infections: Epithelial wounding to cell signaling involved in virus entry. Papillomavirus Res 2019, 7:188–192.

24. Schelhaas M, Shah B, Holzer M, Blattmann P, Kühling L, Day PM, Schiller JT, Helenius A: Entry of human papillomavirus type 16 by actin-dependent, clathrin- and lipid raft-independent endocytosis. PLoS Pathog 2012, 8:e1002657.

25. Bannach C, Brinkert P, Kühling L, Greune L, Schmidt MA, Schelhaas M: Epidermal Growth Factor Receptor and Abl2 Kinase Regulate Distinct Steps of Human Papillomavirus 16 Endocytosis. J Virol 2020, 94.

26. Griffin LM, Cicchini L, Pyeon D: Human papillomavirus infection is inhibited by host autophagy in primary human keratinocytes. Virology 2013, 437:12–19.

27. Surviladze Z, Sterk RT, DeHaro SA, Ozbun MA: Cellular entry of human papillomavirus type 16 involves activation of the phosphatidylinositol 3-kinase/Akt/mTOR pathway and inhibition of autophagy. J Virol 2013, 87:2508– 2517.

28. Ozbun MA, Campos SK: The long and winding road: human papillomavirus entry and subcellular trafficking. Curr Opin Virol 2021, 50:76–86.

29. Smith JL, Campos SK, Wandinger-Ness A, Ozbun MA: Caveolin-1-dependent infectious entry of human papillomavirus type 31 in human keratinocytes proceeds to the endosomal pathway for pH-dependent uncoating. J Virol 2008, 82:9505–9512.

30. Harwood MC, Dupzyk AJ, Inoue T, DiMaio D, Tsai B: p120 catenin recruits HPV to γ-secretase to promote virus infection. PLoS Pathog 2020, 16:e1008946.

31. Inoue T, Zhang P, Zhang W, Goodner-Bingham K, Dupzyk A, DiMaio D, Tsai B: γ-Secretase promotes membrane insertion of the human papillomavirus L2 capsid protein during virus infection. J Cell Biol 2018, 217:3545–3559.

32. Bronnimann MP, Chapman JA, Park CK, Campos SK: A transmembrane domain and GxxxG motifs within L2 are essential for papillomavirus infection. J Virol 2013, 87:464–473.

33. DiGiuseppe S, Keiffer TR, Bienkowska-Haba M, Luszczek W, Guion LGM, Müller M, Sapp M: Topography of the Human Papillomavirus Minor Capsid Protein L2 during Vesicular Trafficking of Infectious Entry. J Virol 2015, 89:10442–10452.

34. Pim D, Broniarczyk J, Bergant M, Playford MP, Banks L: A Novel PDZ Domain Interaction Mediates the Binding between Human Papillomavirus 16 L2 and Sorting Nexin 27 and Modulates Virion Trafficking. J Virol 2015, 89:10145– 10155.

35. Bergant Marušič M, Ozbun MA, Campos SK, Myers MP, Banks L: Human papillomavirus L2 facilitates viral escape from late endosomes via sorting nexin 17. Traffic 2012, 13:455–467.

36. Lipovsky A, Popa A, Pimienta G, Wyler M, Bhan A, Kuruvilla L, Guie M-A, Poffenberger AC, Nelson CDS, Atwood WJ, et al.: Genome-wide siRNA screen identifies the retromer as a cellular entry factor for human papillomavirus. Proc Natl Acad Sci U S A 2013, 110:7452–7457.

37. Popa A, Zhang W, Harrison MS, Goodner K, Kazakov T, Goodwin EC, Lipovsky A, Burd CG, DiMaio D: Direct binding of retromer to human papillomavirus type 16 minor capsid protein L2 mediates endosome exit during viral infection. PLoS Pathog 2015, 11:e1004699.

38. Day PM, Thompson CD, Schowalter RM, Lowy DR, Schiller JT: Identification of a role for the trans-Golgi network in human papillomavirus 16 pseudovirus infection. J Virol 2013, 87:3862–3870.

39. DiGiuseppe S, Luszczek W, Keiffer TR, Bienkowska-Haba M, Guion LGM, Sapp MJ: Incoming human papillomavirus type 16 genome resides in a vesicular compartment throughout mitosis. Proc Natl Acad Sci U S A 2016, 113:6289– 6294.

40. Calton CM, Bronnimann MP, Manson AR, Li S, Chapman JA, Suarez-Berumen M, Williamson TR, Molugu SK, Bernal RA, Campos SK: Translocation of the papillomavirus L2/vDNA complex across the limiting membrane requires the onset of mitosis. PLoS Pathog 2017, 13:e1006200.

41. Aydin I, Villalonga-Planells R, Greune L, Bronnimann MP, Calton CM, Becker M, Lai K-Y, Campos SK, Schmidt MA, Schelhaas M: A central region in the minor capsid protein of papillomaviruses facilitates viral genome tethering and membrane penetration for mitotic nuclear entry. PLoS Pathog 2017, 13:e1006308.

42. Day PM, Baker CC, Lowy DR, Schiller JT: Establishment of papillomavirus infection is enhanced by promyelocytic leukemia protein (PML) expression. Proc Natl Acad Sci U S A 2004, 101:14252–14257.

43. Guion L, Bienkowska-Haba M, DiGiuseppe S, Florin L, Sapp M: PML nuclear body-residing proteins sequentially associate with HPV genome after infectious nuclear delivery. PLoS Pathog 2019, 15:e1007590.

44. Schweiger L, Lelieveld-Fast LA, Mikuličić S, Strunk J, Freitag K, Tenzer S, Clement AM, Florin L: HPV16 Induces Formation of Virus-p62-PML Hybrid Bodies to Enable Infection. Viruses 2022, 14:1478.

45. Longworth MS, Laimins LA: Pathogenesis of human papillomaviruses in differentiating epithelia. Microbiol Mol Biol Rev 2004, 68:362–372.

46. McBride AA: Human papillomaviruses: diversity, infection and host interactions. Nat Rev Microbiol 2022, 20:95–108.

47. van Weering JRT, Verkade P, Cullen PJ: SNX-BAR proteins in phosphoinositide-mediated, tubular-based endosomal sorting. Semin Cell Dev Biol 2010, 21:371–380.

48. Da Graça J, Morel E: Canonical and Non-Canonical Roles of SNX1 and SNX2 in Endosomal Membrane Dynamics. Contact (Thousand Oaks) 2023, 6:25152564231217867.

49. van Weering JRT, Sessions RB, Traer CJ, Kloer DP, Bhatia VK, Stamou D, Carlsson SR, Hurley JH, Cullen PJ: Molecular basis for SNX-BAR-mediated assembly of distinct endosomal sorting tubules. EMBO J 2012, 31:4466–4480.

50. Carlton J, Bujny M, Peter BJ, Oorschot VMJ, Rutherford A, Mellor H, Klumperman J, McMahon HT, Cullen PJ: Sorting nexin-1 mediates tubular endosome-to-TGN transport through coincidence sensing of high-curvature membranes and 3-phosphoinositides. Curr Biol 2004, 14:1791–1800.

51. Kurten RC, Cadena DL, Gill GN: Enhanced degradation of EGF receptors by a sorting nexin, SNX1. Science 1996, 272:1008–1010.

52. Rojas R, Kametaka S, Haft CR, Bonifacino JS: Interchangeable but essential functions of SNX1 and SNX2 in the association of retromer with endosomes and the trafficking of mannose 6-phosphate receptors. Mol Cell Biol 2007, 27:1112–1124.

53. Simonetti B, Daly JL, Cullen PJ: Out of the ESCPE room: Emerging roles of endosomal SNX-BARs in receptor transport and host–pathogen interaction. Traffic 2023, 24:234–250.

54. Simonetti B, Danson CM, Heesom KJ, Cullen PJ: Sequence-dependent cargo recognition by SNX-BARs mediates retromer-independent transport of CI-MPR. J Cell Biol 2017, 216:3695–3712.

55. Kvainickas A, Jimenez-Orgaz A, Nägele H, Hu Z, Dengjel J, Steinberg F: Cargo-selective SNX-BAR proteins mediate retromer trimer independent retrograde transport. Journal of Cell Biology 2017, 216:3677–3693.

56. Lopez-Robles C, Scaramuzza S, Astorga-Simon EN, Ishida M, Williamson CD, Baños-Mateos S, Gil-Carton D, Romero-Durana M, Vidaurrazaga A, Fernandez-Recio J, et al.: Architecture of the ESCPE-1 membrane coat. Nat Struct Mol Biol 2023, 30:958–969.

57. Atwell B, Chen C-Y, Christofferson M, Montfort WR, Schroeder J: Sorting nexin-dependent therapeutic targeting of oncogenic epidermal growth factor receptor. Cancer Gene Ther 2022, doi:10.1038/s41417-022-00541-7.

58. Ho A, Schwarze SR, Mermelstein SJ, Waksman G, Dowdy SF: Synthetic Protein Transduction Domains: Enhanced Transduction Potential in Vitro and in Vivo1. Cancer Research 2001, 61:474–477.

59. Reichelt J, Haase I: Establishment of spontaneously immortalized keratinocyte lines from wild-type and mutant mice. Methods Mol Biol 2010, 585:59–69.

60. Zhang W, Kazakov T, Popa A, DiMaio D: Vesicular trafficking of incoming human papillomavirus 16 to the Golgi apparatus and endoplasmic reticulum requires γ-secretase activity. mBio 2014, 5:e01777–01714.

61. Cardone G, Moyer AL, Cheng N, Thompson CD, Dvoretzky I, Lowy DR, Schiller JT, Steven AC, Buck CB, Trus BL: Maturation of the human papillomavirus 16 capsid. mBio 2014, 5:e01104–01114.

62. Schneider CA, Rasband WS, Eliceiri KW: NIH Image to ImageJ: 25 years of image analysis. Nat Methods 2012, 9:671–675.

63. Measuring cell fluorescence using ImageJ — The Open Lab Book v1.0. [date unknown],

64. Schrödinger, LLC: The PyMOL Molecular Graphics System, Version 1.8. 2015,

65. Thompson JD, Higgins DG, Gibson TJ: CLUSTAL W: improving the sensitivity of progressive multiple sequence alignment through sequence weighting, position-specific gap penalties and weight matrix choice. Nucleic Acids Res 1994, 22:4673–4680.

66. Szklarczyk D, Kirsch R, Koutrouli M, Nastou K, Mehryary F, Hachilif R, Gable AL, Fang T, Doncheva NT, Pyysalo S, et al.: The STRING database in 2023: protein-protein association networks and functional enrichment analyses for any sequenced genome of interest. Nucleic Acids Res 2023, 51:D638–D646.

67. STRING: functional protein association networks. [date unknown],

68. Pyeon D, Pearce SM, Lank SM, Ahlquist P, Lambert PF: Establishment of human papillomavirus infection requires cell cycle progression. PLoS Pathog 2009, 5:e1000318.

69. Campos SK, Chapman JA, Deymier MJ, Bronnimann MP, Ozbun MA: Opposing effects of bacitracin on human papillomavirus type 16 infection: enhancement of binding and entry and inhibition of endosomal penetration. J Virol 2012, 86:4169–4181.

70. Calton CM, Schlegel AM, Chapman JA, Campos SK: Human papillomavirus type 16 does not require cathepsin L or B for infection. J Gen Virol 2013, 94:1865–1869.

71. Galloway CJ, Dean GE, Marsh M, Rudnick G, Mellman I: Acidification of macrophage and fibroblast endocytic vesicles in vitro. Proc Natl Acad Sci U S A 1983, 80:3334–3338.

72. Aydin I, Weber S, Snijder B, Samperio Ventayol P, Kühbacher A, Becker M, Day PM, Schiller JT, Kann M, Pelkmans L, et al.: Large scale RNAi reveals the requirement of nuclear envelope breakdown for nuclear import of human papillomaviruses. PLoS Pathog 2014, 10:e1004162.

73. Barker DF, Campbell AM: The birA gene of Escherichia coli encodes a biotin holoenzyme synthetase. J Mol Biol 1981, 146:451–467.

74. Schatz PJ: Use of peptide libraries to map the substrate specificity of a peptide-modifying enzyme: a 13 residue consensus peptide specifies biotinylation in Escherichia coli. Biotechnology (N Y) 1993, 11:1138–1143.

75. Campos SK: Subcellular Trafficking of the Papillomavirus Genome during Initial Infection: The Remarkable Abilities of Minor Capsid Protein L2. Viruses 2017, 9.

76. Ikegami S, Taguchi T, Ohashi M, Oguro M, Nagano H, Mano Y: Aphidicolin prevents mitotic cell division by interfering with the activity of DNA polymerase-alpha. Nature 1978, 275:458–460.

77. Li S, Bronnimann MP, Williams SJ, Campos SK: Glutathione contributes to efficient post-Golgi trafficking of incoming HPV16 genome. PLoS One 2019, 14:e0225496.

78. Li C: Specific cell cycle synchronization with butyrate and cell cycle analysis. Methods Mol Biol 2011, 761:125–136.

79. Zhang P, Monteiro da Silva G, Deatherage C, Burd C, DiMaio D: Cell-Penetrating Peptide Mediates Intracellular Membrane Passage of Human Papillomavirus L2 Protein to Trigger Retrograde Trafficking. Cell 2018, 174:1465–1476.e13.

80. Nishimura Y, Itoh K: Involvement of SNX1 in regulating EGFR endocytosis in a gefitinib-resistant NSCLC cell lines. Cancer Drug Resist 2019, 2:539–549.

81. Yang Z, Feng Z, Li Z, Teasdale RD: Multifaceted Roles of Retromer in EGFR Trafficking and Signaling Activation. Cells 2022, 11:3358.

82. Day TF, Kallakury BVS, Ross JS, Voronel O, Vaidya S, Sheehan CE, Kasid UN: Dual Targeting of EGFR and IGF1R in the TNFAIP8 Knockdown Non-Small Cell Lung Cancer Cells. Mol Cancer Res 2019, 17:1207–1219.

83. Brinkert P, Krebs L, Ventayol PS, Greune L, Bannach C, Bucher D, Kollasser J, Dersch P, Boulant S, Stradal TEB, et al.: Endocytic vacuole formation by WASH-mediated endocytosis. bioRxiv 2021, doi:10.1101/2021.06.18.448076.

84. Yong X, Zhao L, Deng W, Sun H, Zhou X, Mao L, Hu W, Shen X, Sun Q, Billadeau DD, et al.: Mechanism of cargo recognition by retromer-linked SNX-BAR proteins. PLoS Biol 2020, 18:e3000631.

85. Gokool S, Tattersall D, Seaman MNJ: EHD1 Interacts with Retromer to Stabilize SNX1 Tubules and Facilitate Endosome-to-Golgi Retrieval. Traffic 2007, 8:1873–1886.

86. Xie J, Zhang P, Crite M, Lindsay CV, DiMaio D: Retromer stabilizes transient membrane insertion of L2 capsid protein during retrograde entry of human papillomavirus. Sci Adv 2021, 7:eabh4276.

87. Siddiqa A, Massimi P, Pim D, Broniarczyk J, Banks L: Human Papillomavirus 16 Infection Induces VAP-Dependent Endosomal Tubulation. J Virol 2018, 92.

88. Siddiqa A, Massimi P, Pim D, Banks L: Diverse Papillomavirus Types Induce Endosomal Tubulation. Front Cell Infect Microbiol 2019, 9:175.

89. Da Graça J, Charles J, Djebar M, Alvarez-Valadez K, Botti J, Morel E: A SNX1-SNX2-VAPB partnership regulates endosomal membrane rewiring in response to nutritional stress. Life Sci Alliance 2023, 6:e202201652.

90. Pujals S, Miyamae H, Afonin S, Murayama T, Hirose H, Nakase I, Taniuchi K, Umeda M, Sakamoto K, Ulrich AS, et al.: Curvature engineering: positive membrane curvature induced by epsin N-terminal peptide boosts internalization of octaarginine. ACS Chem Biol 2013, 8:1894–1899.

91. Zakany F, Mándity IM, Varga Z, Panyi G, Nagy P, Kovacs T: Effect of the Lipid Landscape on the Efficacy of Cell-Penetrating Peptides. Cells 2023, 12:1700.

